# Vault Packaging Dynamics: Cargo Loading Stabilizes MVP Conformation for High-Payload Cytokine Delivery in Cancer Therapy

**DOI:** 10.64898/2026.10.27.762310

**Authors:** Qing Lou, Zachary M. Miller, Abby Zhaoqing Wang, Kushal Baranwal, Randall McNally, Z. Hong Zhou, Evan R. Williams, Leonard H. Rome

## Abstract

Recombinant vaults are dynamic protein cages with a large interior volume that can encapsulate therapeutic cargo. Using integrative cryogenic electron microscopy (cryoEM), cryogenic electron tomography (cryoET) and charge-detection mass spectrometry (CDMS), we characterized recombinant empty and interleukin-2 (IL-2)–packaged human vaults produced in yeast to define assembly, packaging heterogeneity and conformational dynamics relevant to optimized cancer and immune therapeutics. CryoEM maps at 2.79 Å (packaged) and 3.16 Å (empty) reveal a D39-symmetric barrel built from 78 major vault protein (MVP) subunits each with an interaction domain (INT) that anchors a cargo on the inner surface at MVP repeats R3– R4. CDMS deconvolution of empty versus packaged particles yields two IL-2–INT vault populations centered near 100 and 400 molecules per vault, with a maximum of ∼660 copies/vault (i.e., many more copies than the 78 available INT anchors), matching cryoET estimates of ∼150–600. Both methods show that highly packaged vaults are enlarged and distorted. Empty particles are conformationally flexible, which promotes initial cargo entry; loading then stabilizes the barrel and condenses cargo, producing vaults that each contain many hundreds of IL-2 molecules. These data establish the structural basis of vault encapsulation and a quantitative framework for engineering vault-based delivery vehicles.

## Introduction

Vaults are abundant ribonucleoprotein particles present in thousands of copies in most eukaryotic cells. Although their precise biological function remains elusive, their composition and architecture have been extensively characterized^1^. These distinctive ovoid particles measure approximately 40 × 40 × 70 nm, ranking among the largest intracellular assemblies. The vault shell is formed by 78 copies of the ∼100 kDa major vault protein (MVP), which assembles in the cytoplasm from a single MVP mRNA on polyribosomes into a bifold-symmetric container exhibiting 39-fold rotational symmetry and perpendicular two-fold axes^2^. Tightly associated with the inner cavity are multiple copies of the high-molecular-mass proteins TEP1 and PARP4, along with one or more vault RNAs (vtRNAs) bound by TEP1. Stoichiometry of these internal components has not been well characterized and may vary across species, tissues, and cellular conditions.

Expression of MVP cDNA alone in heterologous systems such as insect cells or yeast— organisms lacking endogenous vaults—yields empty recombinant vaults structurally indistinguishable from native particles^1,3^. Recombinant vaults have also been produced in mammalian cells such as HEK293, though these may incorporate low levels of endogenous components^4^. The ability to generate milligram quantities of homogeneous empty vaults has enabled extensive engineering for diverse applications, including drug delivery, enzyme bioreactors, vaccines, and mRNA storage^1,5,6^.

A key feature enabling cargo encapsulation is the high-affinity interaction domain (INT) derived from PARP4, which binds a single site on the inner surface of MVP (located primarily at R3 and R4 repeats) with an apparent dissociation constant of 262 nM^7^. INT-tagged fusion proteins spontaneously enter the vault lumen in vitro, exploiting the non-covalent yet highly stable MVP shell and its intrinsic “breathing” dynamics.^5^ This flexibility and the presence of 78 potential INT-binding sites accommodate a wide range of cargos, from peptides and proteins to lipid discs and even AAV particles^4,8^. Molecular dynamics simulations of yeast-produced empty human vaults further support this conformational flexibility^4^.

Previous biophysical studies used scanning transmission electron microscopy (STEM) and gas-phase electrophoretic mobility molecular analysis (GEMMA) to estimate a vault mass at ∼12.9 MDa and demonstrated stoichiometric cargo packaging, albeit with limitations in resolving large heterogeneous particles.^5,9^ Native mass spectrometry (MS) is sensitive and accurate but such large macromolecular complexes are difficult to characterize with standard high resolution MS methods due to spectral complexity associated with sample heterogeneity.^10^ Charge detection mass spectrometry (CDMS) circumvents this problem by weighing individual ions.^11–14^ Masses determined by CDMS have been used to measure protein aggregation, quantitate virus and lipid nanoparticle packaging, and obtain accurate diameters of synthetic nanoparticles in the 100s of nm.^15–19^

Here, we engineered a yeast (*Komagataella phaffii*) strain constitutively expressing human MVP and co-expressing an interleukin-2 (IL2) protein fused to the INT domain (IL2-INT) to produce large quantities of highly purified empty and packaged vaults. We applied electron microscopy (cryoEM), cryogenic electron tomography (cryoET), and CDMS to elucidate vault dynamics, cargo organization, and the mechanistic interplay between packaging and shell stability, with direct implications for optimizing vault-based IL2 delivery in cancer and immune therapy.

## Results

### CryoEM structures of the human recombinant empty and IL2 packaged vault particles

To establish a structural framework for cargo encapsulation, the structure of recombinant empty vault particles (Supplementary Fig. 1) was determined using cryoEM single-particle analysis (Supplementary Fig. 2a). A 3.16 Å reconstruction of the empty vault was produced (Fig. 1a, Supplementary Fig. 3a-c), revealing a barrel-shaped architecture composed of two identical half-vaults, each containing 39 copies of MVP and together forming a D39-symmetric assembly (Fig. 1b). At the distal end of each half-vault, the 39 MVP subunits reorganize into 13 repeating cap groups, each composed of three MVP subunits with distinct conformations (Fig. 1c, Supplementary Fig. 3d-f). The empty vault measures approximately 41.5 nm in diameter (largest waist region R3) and 70 nm in length, consistent with previously reported vault structures^20^. The resulting atomic model, built using AlphaFold3 predictions as starting points and refined in COOT and Phenix, provides a high-resolution structural reference for subsequent analysis of cargo-packaged vaults.

**Fig. 1.**
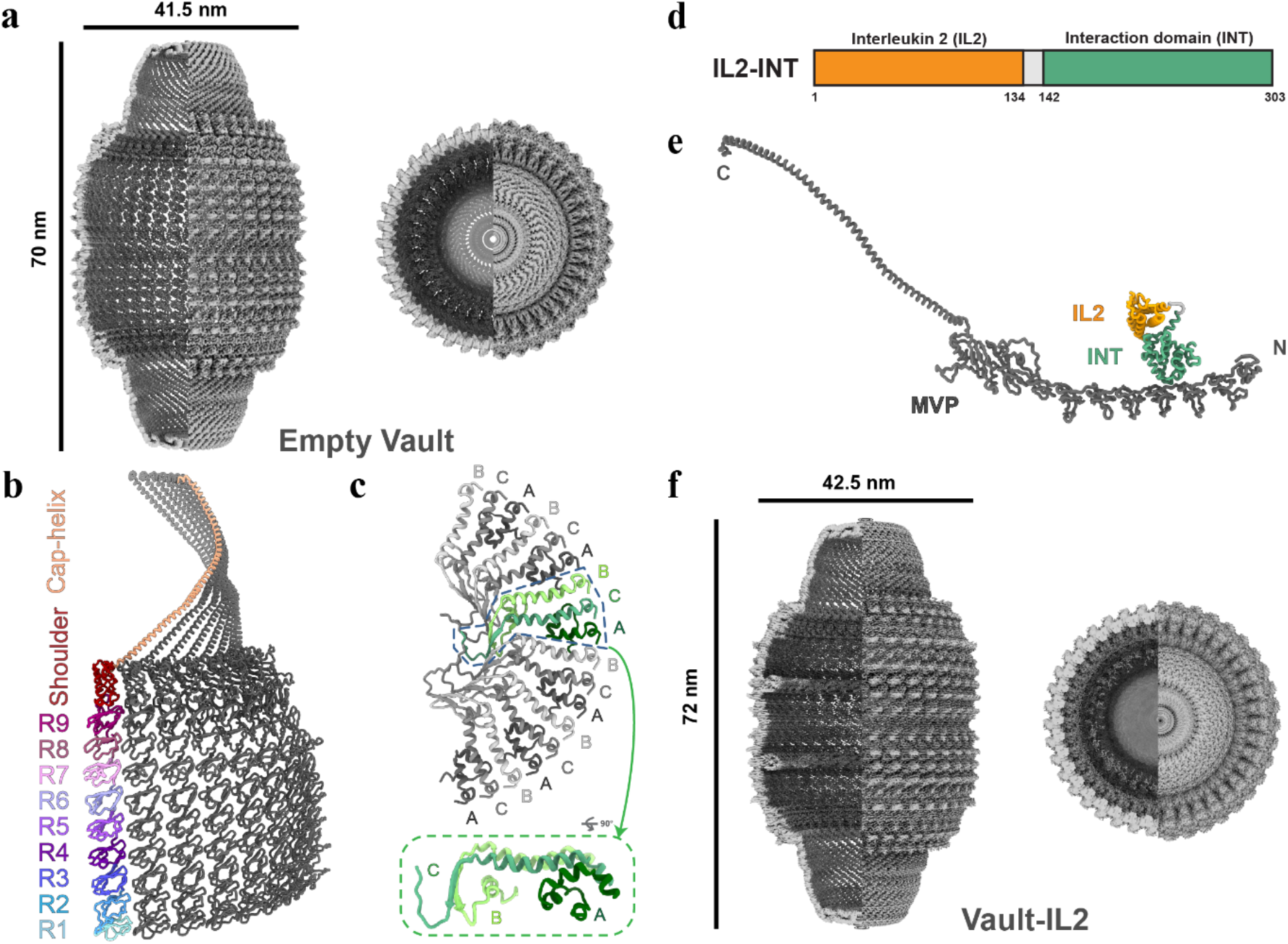
CryoEM structures of the recombinant empty vault and vault–IL2 particles. **a** CryoEM density map of the empty vault (3.16 Å) shown in side and top views. One quarter of the density is clipped away to expose the internal architecture. **b** Arrangement of the 39-fold symmetric portion of the MVP subunits within the vault shell. The displayed model includes residues 2–810. Eight partial MVP subunits are shown, with one highlighted and colored according to its structural domains (R1–R9 repeats, shoulder domain, and cap helix); neighboring subunits are shown in gray. **c** Modeled portion of the vault cap representing the 13-fold symmetry region. The 39 MVP subunits are organized into 13 repeating groups, each containing three MVP subunits in distinct conformations (subunits A, B, and C). Six three-subunit groups are shown; inset shows a zoomed side view of one group (green). **d** Schematic representation of the IL2-INT fusion cargo. The INT domain is fused to IL2 via a 7-amino-acid flexible spacer. **e** Cartoon illustrating the proposed localization of IL2-INT cargo on the inner surface of the vault through INT–MVP interactions. **f** CryoEM density map of the vault–IL2 complex (2.79 Å) shown in side and top views with clipping as in (a). The vault–IL2 particle measures approximately 42.5 nm in diameter and 72 nm in length.

To facilitate cargo encapsulation, interleukin-2 (IL2) was genetically fused to the vault interaction domain (INT) via a flexible GGGGSAS linker (Fig. 1d). This design positions the fused cargo within the vault lumen through high-affinity INT binding to the inner surface of MVP^5^ (Fig. 1e). We then determined the cryoEM structure of vault particles packaged with the IL2-INT cargo (Supplementary Fig. 2b). Single-particle reconstruction yielded a 2.79 Å map of the vault–IL2 complex (Fig. 1f, Supplementary Fig. 3d-f). The overall architecture of the cargo-packaged particle remains highly similar to that of the empty vault, while exhibiting a modest increase in particle dimensions. Additional density was observed within the vault lumen, indicating successful encapsulation of the IL2-INT cargo. These structures establish the architectural basis for subsequent analysis of cargo organization, cargo–vault interactions, and potential conformational changes associated with cargo loading.

### Organization of IL2-INT cargo within the vault

To investigate the spatial organization of the encapsulated cargo, we examined the internal density features within the 2.79 Å cryoEM reconstruction of the vault–IL2 complex. At relatively high contour levels, two prominent concentric density rings are clearly visible inside the lumen, one in each half-vault and located in close proximity to the R3 and R4 repeats of the MVP subunits (Fig. 2a). These rings remain well-defined even at contour thresholds where most of the weaker internal density has disappeared. This persistence indicates a population of cargo molecules that adopt a higher degree of structural order through direct anchoring to the inner surface of the vault shell.

**Fig. 2.**
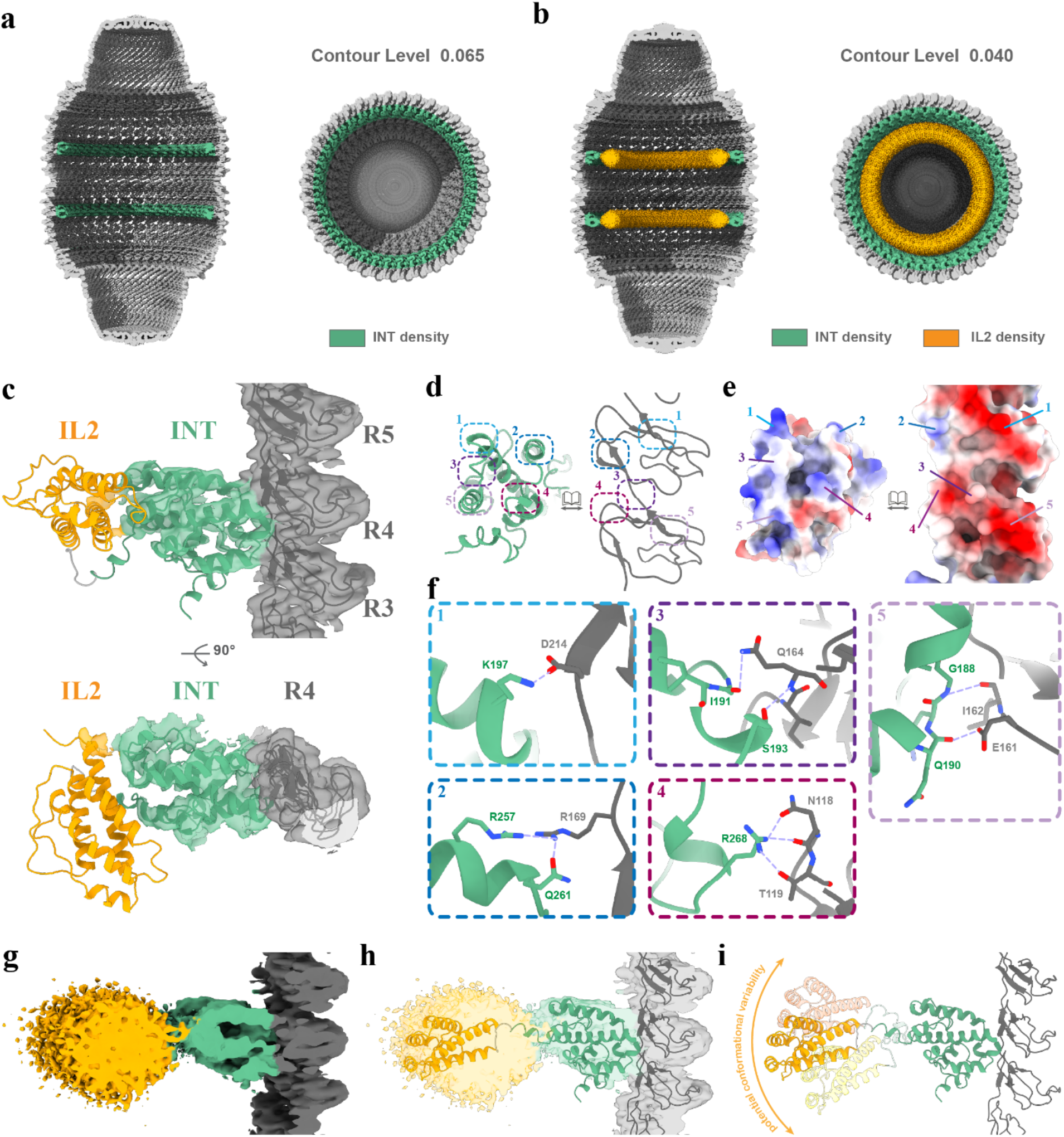
Structural organization and interaction of the encapsulated IL2-INT cargo within the vault. **a** Cut-open side and top views of the vault–IL2 cryoEM density map displayed at a high contour level (0.065). INT domain densities are highlighted in green, revealing two ordered concentric rings. **b** Same views displayed at a lower contour level (0.04), showing additional broader densities (orange) corresponding to IL2. **c** Fitted atomic model of the IL2-INT cargo within the density map. INT (green) anchors the fusion to the inner MVP surface (gray) near R3–R5 repeats; IL2 is shown in orange. **d** Detailed view of the INT–MVP interaction interface. Numbered regions indicate representative interaction sites analyzed in (f). **e** Electrostatic surface of the INT–MVP interface showing complementary charge distributions. **f** Representative intermolecular interactions at the interface (hydrogen bonds and electrostatic contacts with interatomic distances ≤ 3.5 Å). **g** Enlarged view at low contour (0.04) highlighting ordered INT density (green) and disordered IL2 density (orange). **h** Placement of the IL2-INT model within the density shown in (g). **i** Schematic illustration of potential IL2 orientations compatible with the diffuse density, demonstrating conformational flexibility.

Lowering the contour threshold further reveals additional, broader density that extends inward from these ordered rings toward the center of the lumen and forms thicker concentric layers (Fig. 2b). Rigid-body fitting of AlphaFold3-predicted atomic models of the IL2-INT fusion protein showed excellent correspondence for the INT domains within the inner rings, while the IL2 domains accounted for the more diffuse outer density (Fig. 2c). The experimentally observed INT-binding mode at the MVP R3–R4 region aligns closely with the top-ranked solutions from HADDOCK docking, supporting the structural compatibility of the cryoEM-derived INT–MVP interface (Supplementary Fig. 4).

Refinement of the INT–MVP interface identified a network of short-range polar and electrostatic interactions that stabilize anchoring. Key contacts include a putative salt bridge between K197 of INT and D214 of MVP, along with additional hydrogen bonds (Fig. 2d–f). Electrostatic surface potential analysis highlighted strong complementary charge distributions across the binding interface (Fig. 2e). Quantitative assessment using PRODIGY on the refined model yielded a predicted binding free energy of −7.8 kcal/mol, corresponding to a Kd of approximately 2.5 μM at 30 °C. In contrast, the density corresponding to the IL2 portion was markedly broader and less continuous (Fig. 2g), reflecting greater conformational flexibility conferred by the flexible linker. Multiple plausible orientations of IL2 could be accommodated within the diffuse density envelope (Fig. 2h,i), highlighting the high conformational heterogeneity of this distal region. Although the D39-symmetric consensus reconstruction suggests high occupancy of the 78 INT-binding sites as continuous rings, focused 3D classification performed on individual INT-binding regions after symmetry expansion and particle subtraction revealed significant site-to-site variability. Approximately 31% of the classified sites lacked detectable INT density (Supplementary Fig. 5).

We next compared the empty vault (3.16 Å) and IL2–packaged (2.79 Å) structures to evaluate the conformational consequences of cargo anchoring. The two maps share a largely conserved overall architecture, with cargo loading inducing only modest expansion of the vault shell (Supplementary Fig. 6a,b). The largest dimensional increase was observed in the cap region, whereas the shoulder, body, and waist exhibited smaller but measurable changes. Global superposition of individual MVP subunits resulted in an RMSD of 1.21 Å (Supplementary Fig. 6c), indicating that cargo loading causes only minor rearrangements of the MVP fold overall. However, local shifts were evident near the R3–R4 cargo-binding sites when alignments were performed using the cap or waist regions as references.

### Structural and cargo-loading heterogeneity of the vault–IL2 complex

Although the high-resolution consensus reconstruction captured an ordered state of the cargo-packaged vault, it did not fully represent the structural diversity present in the sample. Therefore, we examined the broader particle population through 3D classification and cryoET to assess the full extent of heterogeneity in the vault–IL2 complex. In addition to classes resembling the ordered reconstruction, a substantial fraction of particles exhibited varying degrees of distortion, including expansion of the body/waist region and increased disorder in the cap (Supplementary Fig. 2b). These results indicate that cargo-packaged vaults exist in a continuum of conformational states, with the well-resolved ordered structure representing only one subpopulation within a more diverse ensemble.

To directly visualize structural and cargo-loading heterogeneity in individual particles, we collected cryoET tilt series of the vault–IL-2 sample. Tomogram slices revealed substantial variability in the distribution and density of internal cargo (Fig. 3a). While some vaults displayed relatively compact cargo arrangements adjacent to the shell, others showed extensive density occupying much of the lumen, indicating highly heterogeneous cargo organization. Representative segmentations of individual vault particles further illustrate this diversity, with segmented cargo densities occupying distinct positions and volumes within different particles (Fig. 3b). Such variability is not apparent from the averaged single-particle reconstruction.

**Fig. 3.**
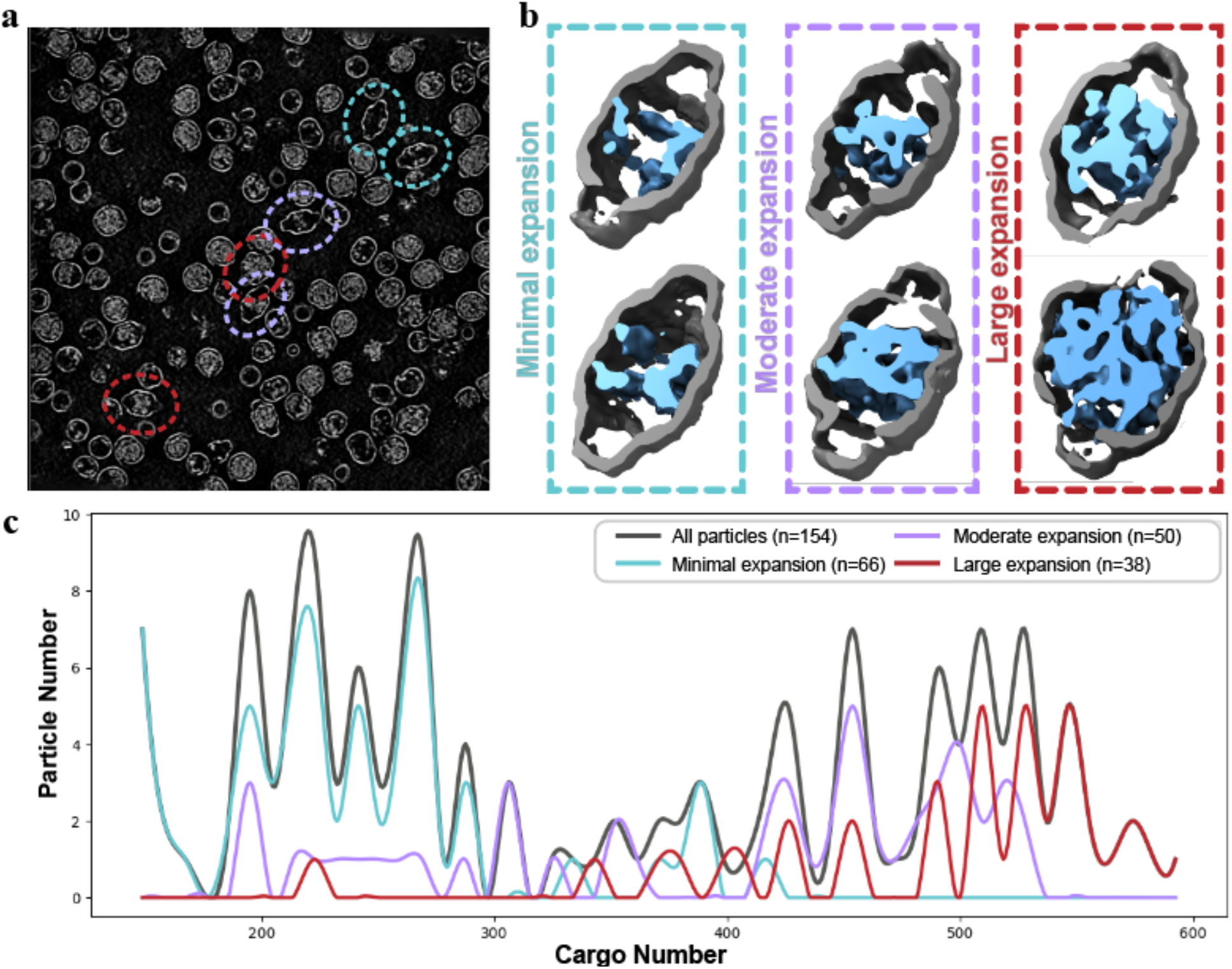
Structural and cargo-loading heterogeneity of vault–IL2 particles revealed by cryo-electron tomography. **a** Representative tomographic slices showing variability in internal cargo density among individual vaults. **b** Segmentations of the particles shown in (**a**), highlighting diversity in cargo organization and occupancy. **c** Distribution of estimated IL2-INT cargo occupancy from cryoET (n = 154 particles). Grey curve shows all particles; colored curves show subsets grouped by transverse expansion (cyan: 1.00–1.15×; purple: 1.15–1.25×; red: 1.25–1.35×).

We then quantitatively examined cargo occupancy across the particles in the tomogram dataset. For each vault, the internal cargo volume was defined as the density enclosed within the vault shell and converted to an estimated number of IL2-INT molecules based on the calculated molecular volume of IL2-INT. The resulting occupancy distribution showed considerable particle-to-particle variability, with estimated cargo numbers ranging from approximately 150 to 600 IL2-INT molecules per vault (Fig. 3c). Notably, these values frequently exceed the 78 shell-associated INT-binding sites resolved in the ordered cryoEM reconstruction, indicating that a substantial portion of cargo is not restricted to the MVP-bound population. To explore the relationship between cargo loading and vault morphology, particles were grouped according to the extent of transverse expansion (measured in the x and y axes). Particles with greater expansion were generally associated with higher cargo occupancies, whereas minimally expanded particles were enriched in lower-occupancy populations. These observations demonstrate the substantial heterogeneity in cargo loading and its correlation with overall vault morphology.

### Cargo loading alters vault structure and dynamics

To more thoroughly characterize the heterogeneity in cargo loading and vault morphology indicated by cryoET, charge detection mass spectrometry (CDMS) was used as an orthogonal single-particle approach. CDMS provides simultaneous measurements of the mass and charge of individual vault particles, allowing both cargo loading and structural heterogeneity to be assessed from the quantitative analysis of thousands of particles. The most abundant peak in the CDMS mass histogram of the unpackaged (empty) vaults (Fig. 4a) is centered at 7.82 MDa and corresponds to intact empty vaults. The measured mass agrees well with the theoretical mass of 7.75 MDa (78 times the 99,327 Da MW of human MVP) but is slightly higher by ∼70 kDa. The 0.9% higher measured mass than theoretical mass and relatively broad peak width are consistent with adduction of non-volatile salts and some remaining solvent typically observed with native mass spectrometry of large macromolecular complexes.^21^ Three low-abundance peaks are centered at 3.87 MDa, 15.72 MDa and ∼11 MDa. These peaks correspond to half-vault particles,^5^ empty vault dimers and vaults containing a broad distribution of unknown cargo, respectively. The latter distribution is likely due to incorporation of proteins and other molecules present during expression of empty vaults.

**Fig. 4.**
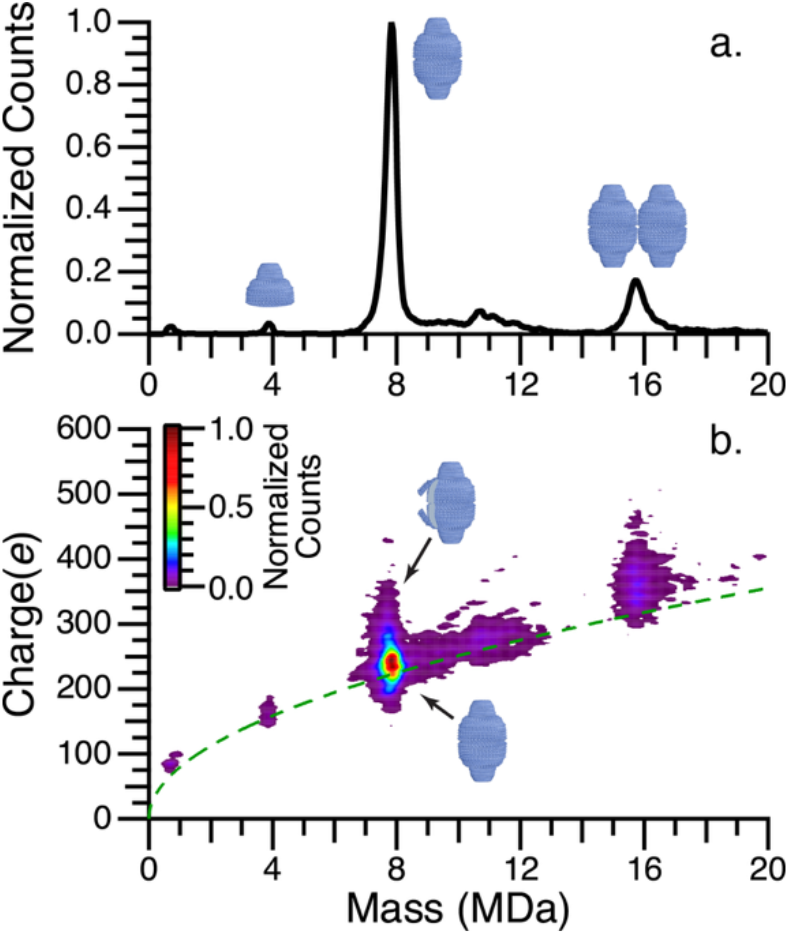
Representative CDMS data of empty vault particles comprised of >6500 ions. **a** 1D mass histogram of empty vault particles. **b** 2D mass / charge histogram of empty vault particles. The Rayleigh limit computed for a spherical water droplet is given by the green dashed line. Vault illustrations are overlaid in (**a**) to connect vault particles with corresponding intact masses and in (**b**) to indicate the regions of the CDMS mass / charge distribution that are attributed to the different structures of the empty vault.

The empty vaults exhibited structural heterogeneity despite their relatively homogeneous mass. The measured charge distribution provides structural information because compact, globular protein complexes typically charge close to but slightly below the Rayleigh limit (see Methods; Equation 1) whereas higher charging indicates more extended or open structures.^22–24^ A 2D mass/charge spectrum of the empty vault particles (Fig. 4b) revealed a remarkably broad charge distribution, peaking at ∼239*e* and ranging from ∼150*e* to ∼450*e*. A small fraction of empty vaults exhibited even higher charge states, extending to greater than 600*e* (Supplementary Fig. 8). Together, these data indicate substantial structural heterogeneity among empty vaults, including a small population of highly extended conformations. The small population suggests that these highly extended structures are transient. This transience may be important for cargo encapsulation by providing access to INT-binding sites on the vault interior.

CDMS data for the IL2-INT packaged vaults were acquired under identical conditions. Mass histograms of the IL2-INT packaged vaults and empty vaults are shown in Fig. 5a (black and blue data, respectively). The packaged vault spectrum contains three distributions of vault particles that are even more apparent in the 2D data (Fig. 5b). The most abundant distribution corresponds to packaged vaults and is centered at ∼11.06 MDa with a full width at half maximum (FWHM) of ∼3.58 MDa. This distribution is ∼3.24 MDa heavier than empty vaults and has a FWHM ∼3.17 MDa greater than that of empty vaults (∼410 kDa). The higher mass and broader peak indicate internalization of ∼93 ± 46 copies of IL2-INT for the main distribution. A second, more highly packaged distribution is centered at ∼23 MDa. A low-abundance tail extending from the mass of the monomer packaged vaults to ∼60 MDa corresponds to vault dimers.

**Fig. 5.**
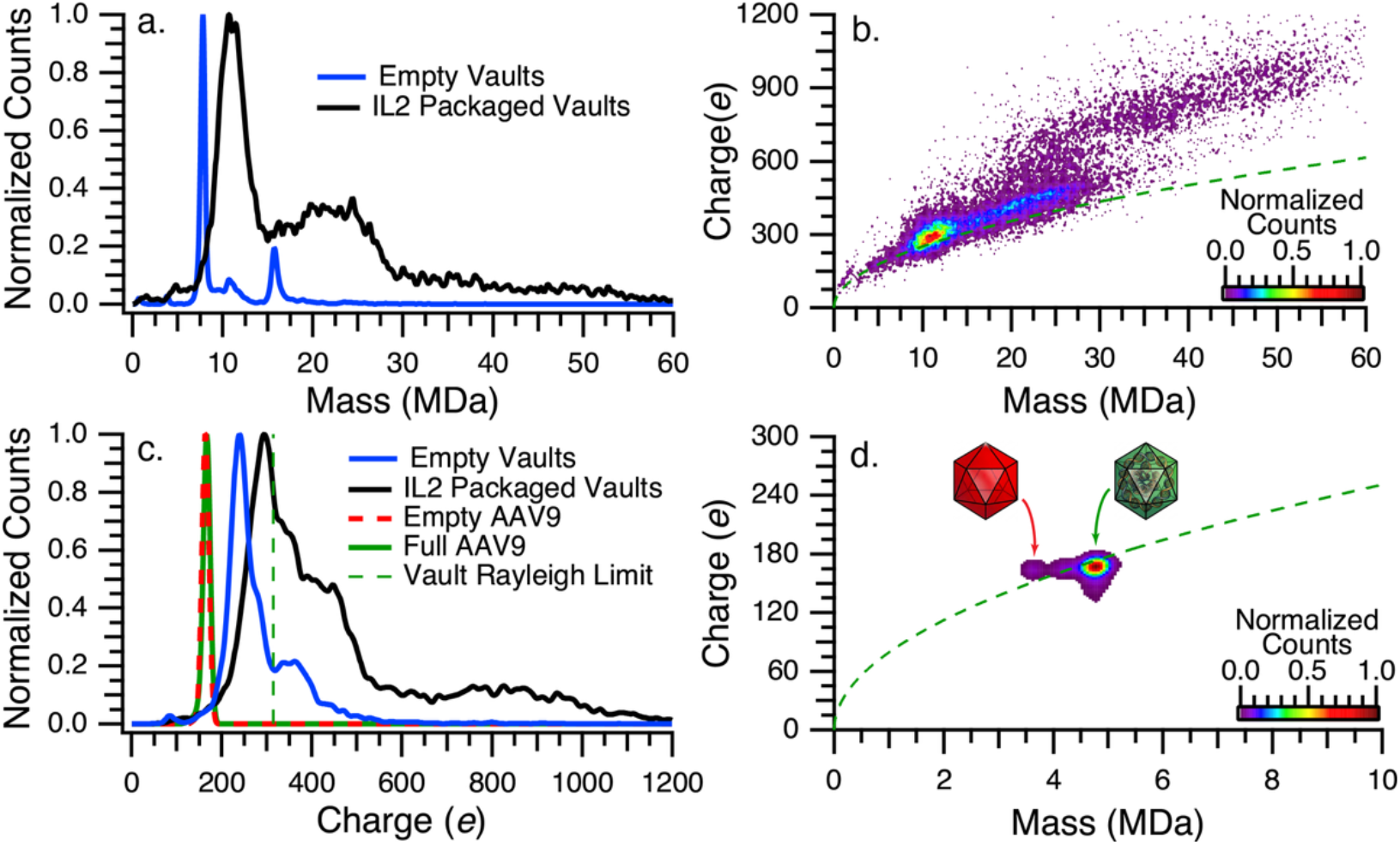
CDMS analysis of IL2-INTpackaged vaults and charge comparison to empty/full AAVs. **a** 1D mass histogram of IL2-INT packaged vault particles (black) and empty vaults (blue). **b** 2D mass / charge histogram of IL2-INTpackaged vaults. **c** Typical 2D mass / charge histogram of AAV9. **d** 1D charge histogram of IL2-INT packaged vaults (black data) and empty vaults (blue data); AAV9 empty capsids (red data) and AAV9 full capsids (green data) are shown for comparison. The estimated Rayleigh limit charging for the vault structure (modeled as an oblate spheroid with an axis ratio of 41.5 nm / 70 nm = 0.59; dimensions shown in Supplementary Fig. S9)^47^ is indicated by the dashed green line in (**c**). The Rayleigh limit for water is given by the green dashed line in (**b**) and (**d**).

The main distribution of monomer packaged vaults centered at ∼11 MDa shows charging slightly above the Rayleigh limit. A second, higher-mass population of monomer packaged vaults extends from ∼15 to ∼28 MDa and exhibits progressively increased charging with increasing IL2-INT cargo mass. This distinct high-mass/high-charge population indicates that extensive cargo loading is accompanied by structural expansion of the vault. This interpretation is supported by the cryoET data (Fig. 3a,b), where vaults with higher internal densities appear distorted and expanded. These bloated packaged vaults account for approximately 45% of the packaged monomer population. A third distribution at overlapping and higher masses (∼20–60 MDa) is attributed to dimers of packaged vaults. This distribution is charged substantially above the Rayleigh limit, consistent with dimers that can retain higher charges than the Rayleigh limit since they typically have non-spherical structures.^14^

Additional insights into the vault structures can be obtained from the charge histograms for packaged and empty vaults shown in Figure 5c (black and blue data, respectively). The predominant packaged-vault population is centered at ∼295*e*, approximately 56*e* higher than the empty-vault population centered at ∼239*e* (Fig. 5c), indicating that the packaged vaults are larger than the empty vaults. The broad distribution in charge states is indicative of structural heterogeneity, although dimers contribute to the high-charge tail. For comparison, empty and DNA-packaged AAV9 capsids, despite their different masses, have nearly identical charge distributions centered at ∼165e (Fig. 5c,d), consistent with well-ordered particles that have essentially the same physical size but different density due to the DNA packaging.^25^

Accounting for the nonspherical shape of the vault gives an estimated Rayleigh limit charging of ∼315e (Fig. 5c, green dashed line). The packaged vaults charge close to this limit (∼94%), consistent with intact vault structures, whereas empty vaults charge substantially below it (∼76%). This difference suggests that empty vaults are more susceptible to compaction, consistent with a more flexible and less structurally constrained shell without cargo present that may undergo gas-phase compaction.^26–28^ These results indicate that the predominant packaged vault population is not distorted but the packaged population at higher masses is structurally expanded.

### CDMS shows cargo loading produces distinct low- and high-occupancy vault populations

The broad mass distribution of the packaged vaults indicated substantial heterogeneity in cargo loading. To quantify the distribution of IL2-INT molecules, the mass distribution of packaged vault monomers was deconvolved using the experimentally measured mass distribution of empty vaults (Fig. 6a,b). The fitted distribution and experimental data are in excellent agreement. Conversion of the deconvolved cargo mass to copy number using the molecular mass of IL2-INT (34.5 kDa) revealed packaging ranging from 0 to ∼660 molecules per vault, with two distinct subpopulations centered at ∼100 and ∼400 copies (Fig. 6c). Remarkably, the bimodal cargo distribution determined by CDMS closely agrees with the range of cargo occupancies independently estimated from the cryoET data (Fig. 3), providing orthogonal evidence for distinct vault populations differing in both cargo loading and physical size. Deconvolution of the packaged-vault dimers yielded a slightly higher average occupancy (∼281 IL2-INT molecules per vault) than observed for monomers (∼230), suggesting that highly packaged, expanded vaults may have an increased propensity to dimerize (Supplementary Fig. 10).

**Fig. 6.**
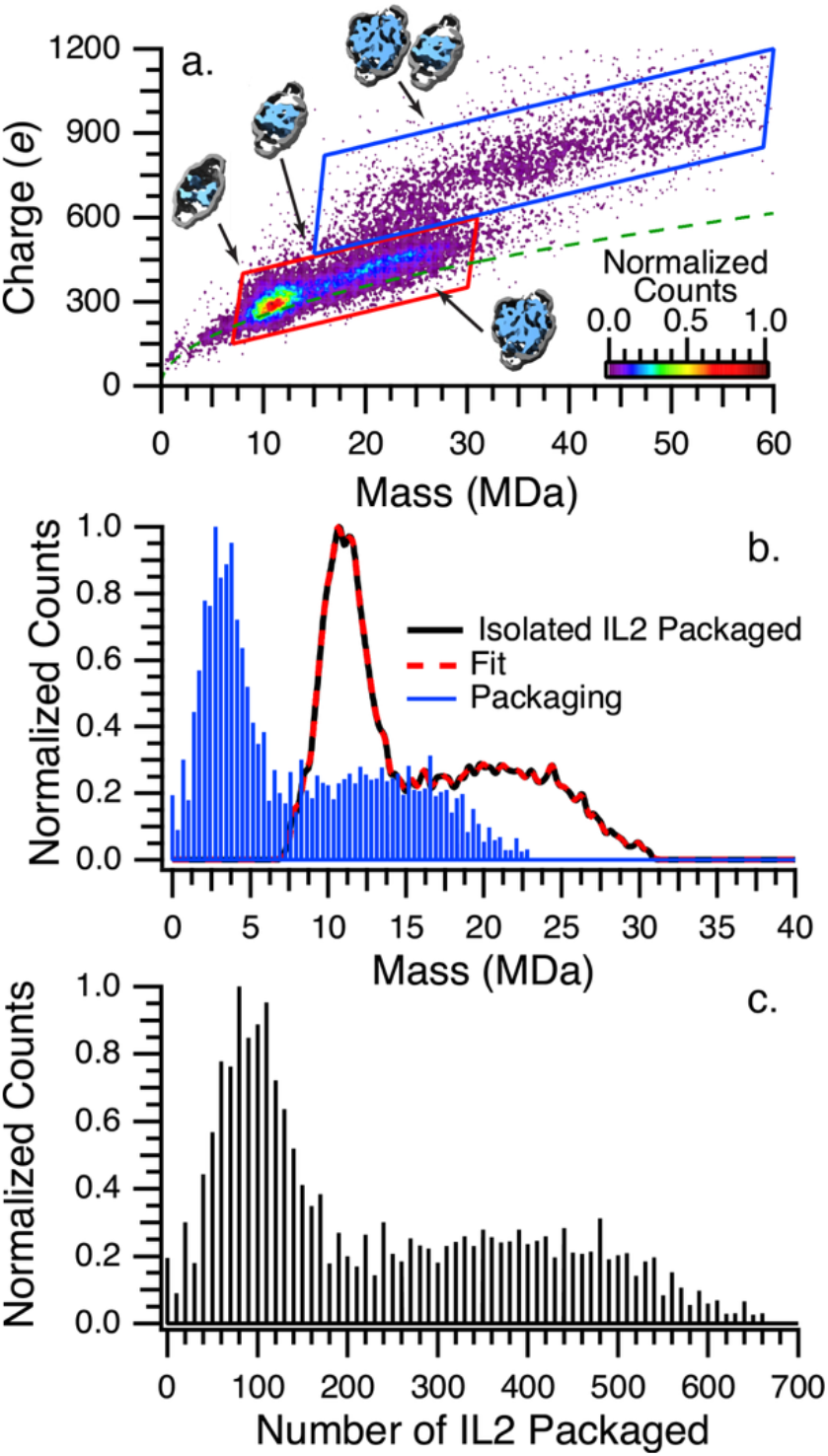
Deconvolution of IL2-INT packaging within vaults. **a** 2D mass / charge histogram of IL2-INT packaged vaults from Figure 5b. Ions within the red and blue polygons are attributed to monomer IL2-INT packaged vaults and dimers of IL2-INT packaged vaults, respectively. The Rayleigh limit for water is given by the green dashed line. IL2-INT vault structures from Figure 3b are overlaid here to indicate areas of the CDMS mass / charge histogram that are attributed to these structures. **b** 1D mass histogram of IL2-INT packaged vault monomers isolated with the red polygon (black data), least squares fit (red dashed line) and the deconvolved packaging mass distribution (blue data). The blue lines are spaced by 10 equivalents of the IL2-INT protein (34.5 kDa). **c** Packaging distribution from (**b**) where the mass axis has been converted to the number of IL2-INT copies.

The bimodal IL2-INT distribution shows that cargo loading is not a continuum but rather comprised of two distinct packaged-vault populations. The lower-occupancy population, centered at ∼100 IL2-INT copies, corresponds primarily to vaults that retain their native dimensions. This occupancy exceeds the 78 known INT-binding sites, indicating that additional IL2-INT molecules can be retained within the vault without direct INT–MVP binding. The higher-occupancy population consists of expanded vaults containing an average of ∼400 IL2-INT molecules. Based on the internal volume of an undistorted vault (3.87×10^−20^ liters; Supplementary Fig. 9) and the approximate density of protein (∼1.42 g/mL),^29^ this corresponds to an effective concentration of ∼17 mM for vaults containing 400 copies of IL2-INT, indicating that the cargo is densely packed within the confined vault lumen. Such high local concentrations are likely to promote cargo–cargo interactions and may drive formation of a nano-condensate or an amorphous solid that can induce nanocrystal formation.^30^ Formation of this densely packed state is accompanied by vault deformation, which ultimately means that the integrity of the vault structure limits the maximum cargo that can be accommodated simply due to the internal volume of a well-formed empty vault versus a distorted vault.

## Discussion

Our integrative cryoEM, cryoET, and CDMS analyses provide the first high-resolution view of recombinant human vaults in both empty and densely cargo-packaged states. The atomic structures reveal how INT-tagged IL2 is anchored at the R3–R4 repeats of MVP while maintaining overall vault architecture with only modest shell expansion. The combination of well-ordered INT rings and diffuse IL2-INT density, together with direct visualization of particle-to-particle variability by cryoET and corresponding mass shifts by CDMS, demonstrates that cargo loading is inherently heterogeneous. Importantly, the data show that the majority of vaults have IL2-INT cargo that exceeds the 78 canonical INT-binding sites, indicating that the large internal volume serves as a reservoir for additional molecules that are dynamically exchanging with shell-associated sites. The high effective concentration of IL2-INT inside the vaults can lead to the initiation of nano-condensate or crystallization of IL2-INT molecules within the vaults leading to a significant population of highly packaged vaults that have distorted shapes to accommodate the large number of IL2-INT molecules that are contained.

The bimodal IL2-INT cargo distribution could be explained by nano-condensate formation of IL2-INT within the vaults. The lower packaging distribution centered ∼100 IL2-INT may correspond to a steady-state in which IL-2 is initially trapped inside the vault by a significant fraction of the 78 known binding sites. As more IL2-INT enters the vaults, close interactions with IL2-INT or the protein interior leads to a higher average packing compared to the known 78 binding sites, a process that may be enhanced by reduced flexibility of the packaged vaults indicated in both the CDMS and the cryo-EM data. Conversion of these structures to the more highly packaged bloated structures may require a critical mass of IL2-INT that initiates nano-condensate formation. In some fraction of vaults that reach this critical mass, rapid condensation may occur, allowing additional IL2-INT to enter the vault and condense. Condensation processes are typically rapid, especially in confined spaces with high protein concentration which leads to the observed bimodal packaging distribution. The nano-condensation process is ultimately retarded by the confinement within the vault, although this appears to also lead to bloated vaults. The nano-condensation process is likely reversible given the small size. This mechanism suggests that the relative abundances of the two packing populations will likely be sensitive to packaging conditions.

A central mechanistic insight from this work is that cargo loading actively modulates vault dynamics. Comparison of empty and IL2–packaged structures, supported by improved local resolution in the body region upon packaging, indicates that INT–MVP interactions stabilize the inward-facing conformation of MVP subunits. This stabilization reduces “breathing” motions of the shell, creating a self-reinforcing state in which higher cargo occupancy further constrains flexibility. The CDMS charge distributions and cryoET occupancy measurements are fully consistent with this model: empty vaults are conformationally heterogeneous with a fraction of the high-charge-state population indicating more open structures. This bidirectional relationship — where the vault’s flexibility enables cargo entry, which in turn restricts flexibility and traps cargos within — explains why vault packaged therapeutics do not leak out of the particle and supports the need for cargo release to be triggered by external forces, such as low pH of the endosome or proteolytic activity.

The fish-trap analogy captures the gist of the above mechanism of this process (Fig. 7). Transient openings of individual MVP chains expose INT-binding sites, allowing abundant IL2–INT molecules to enter and bind. As more sites become occupied, the shell becomes progressively more rigid, shifting the equilibrium toward the packaged state and trapping additional cargo within the large internal volume. This mechanism transforms the vault from a passive container into an active, occupancy-dependent trap. This model is consistent with earlier in vitro loading experiments showing that INT-fused cargos can enter preassembled recombinant vaults in a cell-free system. Thus, although MVP and IL2-INT were co-expressed in the yeast system used here, the observed cargo packaging does not need to be explained by co-assembly during vault formation.

**Fig. 7.**
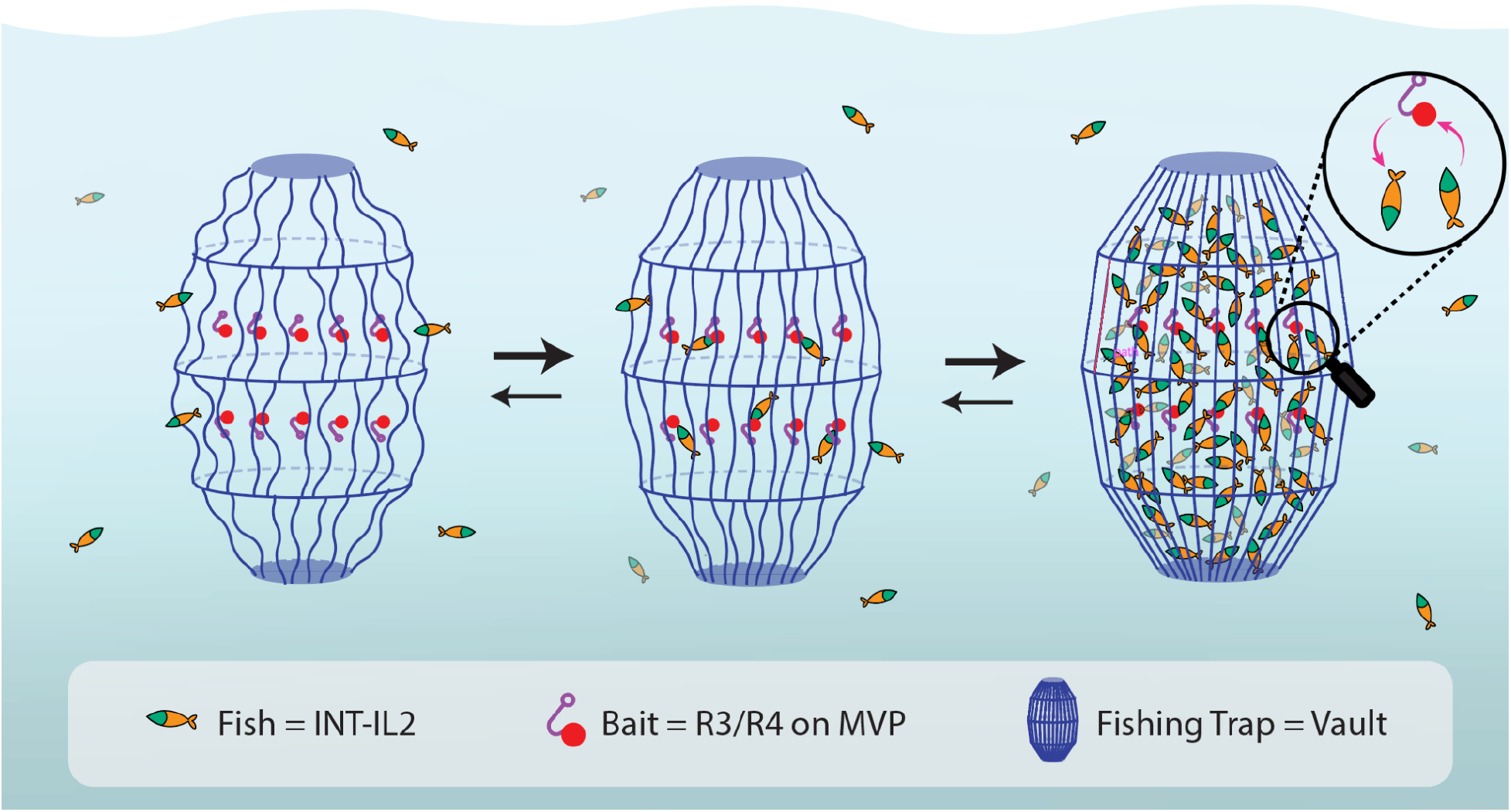
Schematic illustration of our proposed mechanism accounting for loading more than 78 IL2-INT molecules/vault by analogy to a fish trap. The vault shell is represented as a dynamic trap, IL2-INT cargos as fish, and MVP R3/R4 repeats as bait. In the low-occupancy (empty) state, the shell is flexible (wavy lines). As cargo accumulates, the shell becomes more ordered and rigid (straighter lines). The fully packaged state shows dense, heterogeneous internal cargo. Inset highlights the dynamic binding equilibrium at individual MVP sites.

From a translational perspective, these findings have immediate implications for vault-based therapeutics. The ability to achieve high-density packaging of potent cargos such as IL2 while maintaining particle stability addresses a long-standing challenge in nanoparticle drug delivery. The self-reinforcing packaged state ensures efficient encapsulation and protection during circulation, yet the inherent dynamics of the vault shell suggest that controlled release can be triggered by environmental cues (e.g., low pH in endosomes or proteolytic processing), offering a “Goldilocks” solution — high packing density combined with tunable release.

These findings may also provide insight into how endogenous vault-associated proteins are accommodated within native vault particles. In this study, high-density cargo packaging was achieved by fusing IL2 to the INT domain, which mimics the vault-interaction region of PARP4. In native vaults, however, PARP4 and TEP1 are substantially larger than IL2-INT. Using the INT-bound position observed in our vault–IL2 structure as a guide, manual steric-feasibility modeling (Supplementary Fig. 11a) suggests that multiple PARP4 molecules could be accommodated within the vault lumen without major steric overlap, with an approximate upper range of 6-7 copies per half vault depending on conformation (Supplementary Fig. 11c). Similar modeling of TEP1, positioned with its N-terminal/vtRNA-associated region oriented toward the cap-proximal lumen, suggests that ∼2-3 TEP1 molecules per half vault could also be accommodated (Supplementary Fig. 11b), although the precise number and orientation are likely to depend on flexible regions and the presence of vtRNA. As natural vaults are highly stable and long-lived particles,^31^ it is likely that they reach a packaging stoichiometry that is dependent on the cellular concentrations of PARP4 and TEP1 and these vaults will be fully packaged and thus conformationally restrained, implying that vault content proteins do not freely exchange between the vault interior and the cytoplasm. Concentrations of PARP4 and TEP1 are likely to vary in different cells and tissues, leading to structural and functional heterogeneity.

Collectively, our results establish recombinant vaults as a highly optimized platform to package cargo molecules, such as IL2 for cancer therapy. By harnessing the natural breathing of the vault shell, feedback of cargo loading, and high-density packaging enabled by loading enhanced cargo condensation, we can reliably encapsulate hundreds of therapeutic molecules per particle while preserving the particle’s primary structural integrity and targeting potential. This work opens the door to next-generation vault-based nano-medicines that combine high payload capacity, stability in circulation, and environmentally triggered release, bringing us significantly closer to safe and effective delivery of potent immunotherapeutics such as IL2 directly to tumor sites.

## Methods

### Cloning, expression, and purification of vaults

Codon-optimized DNA encoding human MVP was inserted into vector pJGG (BioGrammatics, San Diego, CA) and transformed into yeast (*Komagataella phaffii,* previously called *Pichia pastoris*) cells. MVP-expressing *Pichia* was cultured in YPD medium at 30 °C in shake flasks at 200 rpm for 24 hours. Harvested cells were resuspended in Buffer A (50 mM Tris pH 7.4, 75 mM NaCl, 0.5 mM MgCl₂) supplemented with 1 mM DTT and protease inhibitor cocktail (Sigma #P8215) and lysed using an Avestin Emulsiflex C3 homogenizer. Cellular debris was removed by centrifugation at 20,000 × g for 20 minutes. Supernatant was further centrifuged at 37,000 rpm for 1 hour, and the vault-containing pellet was resuspended in Buffer A containing 1% Triton X-100. Vaults were precipitated by addition of 0.1 g ammonium acetate per mL and agitated overnight at 4 °C. Precipitate was collected by centrifugation at 20,000 × g for 20 minutes, resuspended in Buffer A, and remaining debris was removed by centrifugation at 3,000 × g for 10 minutes. Supernatant was applied to a TMAE Fractogel HiCap column (EMD) and eluted via NaCl gradient. Vault-containing fractions were pelleted at 37,000 rpm for 1 hour and resuspended to a concentration of 1 mg/mL in phosphate-buffered saline containing 10 mg/mL trehalose (PBS). Aliquots of resuspended vaults were frozen in liquid nitrogen, or lyophilized and stored at –20 °C^1,3^. Studies show that packaged vaults are stable for months stored at 4 °C in PBS and do not lose packaged cargo.

Sequences encoding human IL2-INT fusion proteins, separated by GGGGSAS spacers, were separately inserted into vector pJUH (BioGrammatics) and transformed into MVP-expressing *Pichia* cells. Colonies expressing MVP and IL2-INT were selected. The resulting IL2-INT-containing vaults were expressed and purified as above.

Purified empty vault and vault–IL2 samples were characterized by SDS-PAGE with Coomassie blue staining to assess protein composition, negative-stain EM to assess particle morphology, and quantitative ELISA to estimate MVP and IL2-INT content in vault–IL2 particles (Supplementary Fig. 1).

### CryoEM sample preparation and image acquisition

For vitrification of both empty vault and vault-IL2 samples, 3 μL of specimen was applied onto glow-discharged Quantifoil holey carbon grids (200 mesh, R2/1). Grids were glow-discharged for 90 s at 25 mA by using a PELCO easiGlow system prior to sample application. Excess liquid on the grids was removed using a Vitrobot Mark IV (FEI/Thermo Fisher) operated at 4°C and 100% humidity with a blot force of 2 and a blot time of 5 s. The grids were subsequently plunge-frozen in a precooled liquid ethane/propane mixture. Grid quality, plunge-freezing conditions, and sample concentration were optimized through screening on an FEI TF20 transmission electron microscope equipped with a Gatan K2 camera.

Single particle cryoEM data for empty vault specimen were collected as movie stacks on optimized grids using an FEI Titan Krios G1 (300 kV) equipped with an energy filter (20 eV slit width) and a Gatan K3 camera operated at super-resolution mode (81,000×, 0.55 Å/pixel). Automated data collection for the empty vault dataset was performed using SerialEM^32^. For vault-IL2 sample, cryoEM data were collected on an FEI Titan Krios G4 (300 kV) with an energy filter (10 eV slit width) and a Falcon 4i camera (130,000×, 0.97 Å/pixel). Automated data collection for the empty vault dataset was performed using SerialEM. For the vault–IL2 sample, cryoEM data were collected on an FEI Titan Krios G4 microscope operating at 300 kV and equipped with an energy filter with a 10 eV slit width and a Falcon 4i camera operated at a nominal magnification of 130,000×, corresponding to a pixel size of 0.97 Å/pixel. Automated data collection for the vault–IL2 dataset was performed using EPU. Each movie stack was fractionated into 45 frames with a total accumulated dose of 50 e/Å²/micrograph. Data were collected with a defocus range of −1.5 to −3.0 μm.

### CryoEM data processing and model building

Motion correction and dose weighting of all movie frames were performed using MotionCor2^33^. A total of 26,966 and 32,192 drift-corrected micrographs for empty vault and vault-IL2 datasets, respectively, were imported into cryoSPARC^34^ for subsequent image processing. For the reconstruction of empty vault structure, 3,920,582 particles were picked using Template Picker with templates generated from a previously reported human vault density map (EMD-53805). Following 2D classification, 407,778 particles were selected for Ab-Initio Reconstruction and Homogenous Refinement with D39 symmetry applied using bin-4 particles. The aligned particle coordinates were subsequently re-extracted without binning and subjected to 3D classification without alignment or symmetry. A subset of 126,007 particles (30.9%) from classes exhibiting minimal structural distortion were selected for final reconstruction. Homogeneous refinement with defocus and global contrast transfer function (CTF) refinement under D39 symmetry yielded a density map at an overall resolution of 3.16 Å.

For focused analysis of the vault cap region, particles from the empty-vault dataset used for the D39-symmetric reconstruction were subjected to D1 symmetry expansion to separate the two symmetry-related cap regions. The resulting particles were re-extracted with the cap region centered in the box, yielding 251,981 cap-centered particles. Particle subtraction was then performed using a soft cap mask to retain the cap density and remove signal from the remaining vault body. The cap-centered, subtracted particles were subjected to C1 3D classification. Particles from classes with well-defined cap density were selected for final refinement, resulting in 164,854 particles. Refinement with C13 symmetry yielded a cap-focused reconstruction at 3.12 Å resolution.

For the reconstruction of vault-IL2, 382,621 particles were picked using Blob Picker with particle diameters ranging from 400 to 900 Å. Following 2D classification, 318,865 particles were selected for Ab-Initio Reconstruction and Homogenous Refinement with D39 symmetry applied using bin-3 particles. The aligned particle coordinates were subsequently re-extracted without binning and subjected to 3D classification without alignment. A subset of 69,510 particles (21.8%) from classes without distortion were selected and further reconstructed into 2.79 Å density map using Homogeneous refinement with defocus and global CTF refinement under D39 symmetry.

To examine occupancy of the shell-associated IL2-INT cargo, focused 3D classification was performed on the INT-binding region of the vault–IL2 reconstruction. Particles from the D39-symmetric vault–IL2 reconstruction were subjected to symmetry expansion to generate particle copies corresponding to symmetry-related INT-binding sites. A soft mask encompassing the local INT-binding region, including the INT density and adjacent MVP repeats near R3–R4, was used for particle subtraction to remove signal from the remaining vault shell. The resulting focused particles were classified in 3D without imposing symmetry. Classes were inspected for the presence or absence of INT density at the binding site.

Model building for both structures was initiated using AlphaFold3 predictions.^35^ For the empty vault structure, the MVP model was initially rigid-body fitted into the map of empty vault (3.16 Å) using UCSF ChimeraX^36^. The model was subsequently adjusted manually in Coot^37^ and further refined by real-space refinement in Phenix^38^. For the cap-focused reconstruction, coordinates from PDB: 9R87^39^ were used as an initial model and rigid-body fitted into the cap map (3.12 Å) using UCSF ChimeraX. The cap model was subsequently manually rebuilt in Coot according to the experimental density, followed by real-space refinement in Phenix. For the structure of vault-IL2, the refined MVP model from empty vault structure together with the AlphaFold3 prediction of the IL2-INT complex were rigid-body fitted into the vault-IL2 map (2.79 Å). The MVP and INT domain were manually adjusted in Coot, and the MVP model was further refined by real-space refinement in Phenix. All visualizations of models and maps in figures were prepared using UCSF ChimeraX.

### Computational model analysis and volume estimation

HADDOCK^40^ docking was performed to evaluate whether the IL2-INT cargo model independently supported the INT-binding site observed in the cryoEM reconstruction. The AlphaFold3-predicted IL2-INT model was docked against the MVP subunit, and the top-ranked docking solutions were inspected for the location and orientation of the INT-MVP interaction. The predicted docking solutions were compared with the INT-MVP interface observed in the cryoEM-derived model.

The INT-MVP interface in the refined model was further analyzed using PRODIGY^41^ to estimate the binding affinity and identify interface properties. The predicted binding affinity and dissociation constant were used as computational support for formation of the INT-MVP interface observed in the reconstruction.

The molecular volume of the IL2-INT cargo was estimated from the AlphaFold3-predicted IL2-INT model in ChimeraX by generating a solvent-excluded molecular surface and measuring the enclosed surface volume as detailed in Supplementary Figure 9. The measured volume was 40.2 nm³ per IL2-INT molecule. The internal vault volume was estimated geometrically. Internal dimensions were measured from the empty-vault structure in ChimeraX, and the vault lumen was approximated as a series of truncated cones. For each segment, the vertical height was calculated from the measured slanted side length and the difference in internal diameters, and the segment volume was calculated using the truncated-cone formula. Segment volumes were summed for one half-vault and doubled to estimate the full vault lumen volume. This calculation gave an approximate full internal volume of 4.25 × 10⁴ nm³, or 3.87 × 10⁴ nm³ when the cap-proximal frustum was excluded. Theoretical cargo capacity was estimated by dividing the vault internal volume by the IL2-INT molecular volume and should be interpreted as a no-solvent upper-bound estimate.

### CryoET tilt series acquisition and data processing

Tilt series were collected using Tomography 5 in an FEI Titan Krios G4 (300 kV) equipped with an energy filter (10 eV slit width) and a Falcon 4i camera. Data was collected at magnification 64,000× (2.00 Å/pixel). Tilt series were collected from −60° to 60° with a 3° tilt increment using a dose-symmetry scheme. The defocus was set from −3.5 μm. Tilt series alignment and tomographic reconstruction were performed using CryoFlow. IsoNet^42^ was applied to enhance tomogram contrast and reduce the missing wedge artifacts. Visualization of tomogram slices and three-dimensional rendering of tomograms were performed using IMOD^43^ and UCSF ChimeraX.

### Quantification of packaged IL2-INT molecules within individual vault particles

Analysis was performed by first generating spatial fits for vault particles identified in four tomograms. Vaults judged to be of sufficient quality were manually selected, and local registration was performed using an idealized vault structure derived from empty vault structure. Local registration determined the spatial transformation required to align the idealized model to each selected vault within the tomographic volume. In addition to rotational and translational alignment, isotropic scaling was applied along the x- and y-axes of the model reference frame, corresponding primarily to changes in the vault waist diameter. A total of 154 vault fits were generated across all tomograms.

To account for variation in image contrast across sample depth due to differences in the contrast transfer function (CTF), intensity thresholds were determined independently for each z-slice of every tomogram (Supplementary Fig. 7). Threshold optimization was based on binary classification of voxels into positive and negative classes. The positive class consisted of voxels located within 10 Å of the fitted vault surface, which were expected to correspond primarily to major vault protein (MVP) density. The negative class consisted of voxels located 40–50 Å outside the fitted structure after masking the vault interior to avoid contributions from putative IL2 density. For each z-slice, 800 candidate thresholds were evaluated and the corresponding Matthew’s Correlation Coefficient (MCC) values were calculated. The threshold yielding the maximum MCC value was selected as the optimal threshold. Across all z-slices, the mean MCC value was approximately 0.5.

The optimized thresholds were subsequently used for binary classification of voxels within each fitted vault. Analysis was restricted to the vault waist region, while the cap regions were excluded by masking. Additionally, voxels within 20 Å of the vault wall were omitted to minimize artifacts arising from imperfect fitting accuracy. Percentage voxel occupancy was then calculated as the ratio of the number of voxels within the vault waist exceeding the selected threshold to the total number of voxels in the analyzed interior region. To estimate molecular occupancy, the entire procedure, including fitting, threshold determination, and occupancy calculations, was repeated using an atomic model converted to a density map with the molmap function in UCSF ChimeraX. This reference density yielded an occupancy of approximately 0.24% for a single IL2-INTbinding domain within a non-scaled vault and was subsequently used to estimate the number of IL2-INT molecules associated with each analyzed vault particle, accounting for the scaling factor.

### Charge detection mass spectrometry

Experiments were performed with a custom-built CDMS instrument that is described in detail elsewhere.^14^ CMV-GFP AAV9 samples were purchased from the University of North Carolina Vector Core. Vault and AAV9 samples were buffer exchanged into 200 mM and 500 mM ammonium acetate, respectively, prior to CDMS measurements. Analyte ions are formed with nanoelectrospray using borosilicate emitters with tip diameters of ∼1.8 µm. Ionized vault particles were trapped for 500 ms, and their induced signals were amplified by a charge-sensitive preamplifier (Amptek A250 CoolFET). Time-domain transients, sampled at 1 MHz with 12 bit precision, are analyzed using a program written in the C++ programming language that performs rectangularly apodized short-time Fourier transforms (STFT) to determine the masses, energies, and charges of the individual ions that are simultaneously trapped.^44,45^ An STFT with a segment length of 25 ms and step size of 5 ms was used in all experiments.

To obtain information about the structures of vaults in solution, the individual ion charge states that were measured were compared to the Rayleigh limit for a spherical water droplet. This limit indicates the theoretical maximum charge state that a stable spherical droplet can have based upon density and surface tension.^46^ Most globular proteins and protein complexes charge near or below the Rayleigh limit.^22–24^ The Rayleigh limit charge (*z_R_*) of a spherical water droplet is given in Equation (1), where *e_0_* is the permittivity of the surrounding media, and *γ* is the surface tension of water, *r* is the radius of the droplet, *e* is the number of elementary charges in one coulomb, *m* is the mass of the droplet, *r* is the density of water and *N_A_* is Avogadro’s constant.

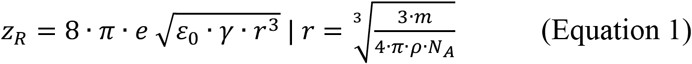

CDMS mass histograms of packaged vaults were fit using a discrete convolution method that has been described previously.^17^ In brief, the mass histogram of empty vaults and a synthesized mass distribution for packaging (here defined by a series of summed Gaussian distributions that are spaced by the mass of 10 equivalents of IL2-INT) are convolved to fit the mass histogram of IL-2 packaged vault monomers. The height of each individual Gaussian was optimized using non-linear least squares to fit the convolved distribution to the mass histogram of packaged vaults.

## Acknowledgements

This research was supported in part by NIH (R01GM071940 to ZHZ and 5R01GM139338 to ERW) and a shared instrumentation CTSI grant to LHR (NIH/National Center for Advancing Translational Science UCLA CTSI Grant UL1TR001881). We acknowledge the use of resources at the Electron Imaging Center for Nanomachines [supported by UCLA and by instrumentation grants from the NIH (1S10OD018111) and NSF (DBI-1338135)]. We thank Professor Joseph A. Loo for sample handling and helpful discussions.

## Author Contributions

Z.H.Z. and L.H.R. conceived the project; Z.H.Z. supervised trainee research performed by Q.L., A.Z.W. and K.B.; R.M. prepared the samples and quantitated sample purity; L.H.R. and Q.L. carried out negative staining; Q.L. recorded cryoEM images and cryoET tilt series and determined the 3D structures; Q.L. and A.Z.W. built the atomic models; Z.H.Z., Q.L., K.B. and A.Z.W. analyzed the cryoET data; Z.M.M. acquired CDMS data; E.R.W and Z.M.M. analyzed and interpreted the CDMS data; Z.H.Z., L.H.R., Q.L., E.R.W., and Z.M.M. wrote the manuscript; and all authors contributed to the editing of the manuscript.

## Accession numbers

The cryoEM density maps and corresponding atomic models have been deposited in the EMDB and PDB, respectively. The accession numbers are listed as follows: empty vault (PDB: XXXX and EMD-XXXX); vault with cargo (PDB: XXXX and EMD-XXXX); cryoET tomograms (EMD-XXXXX and EMD-XXXXX, for empty and cargo-packaged vaults, respectively.

## SUPPLEMENTARY INFORMATION

**Supplementary Fig. 1.**
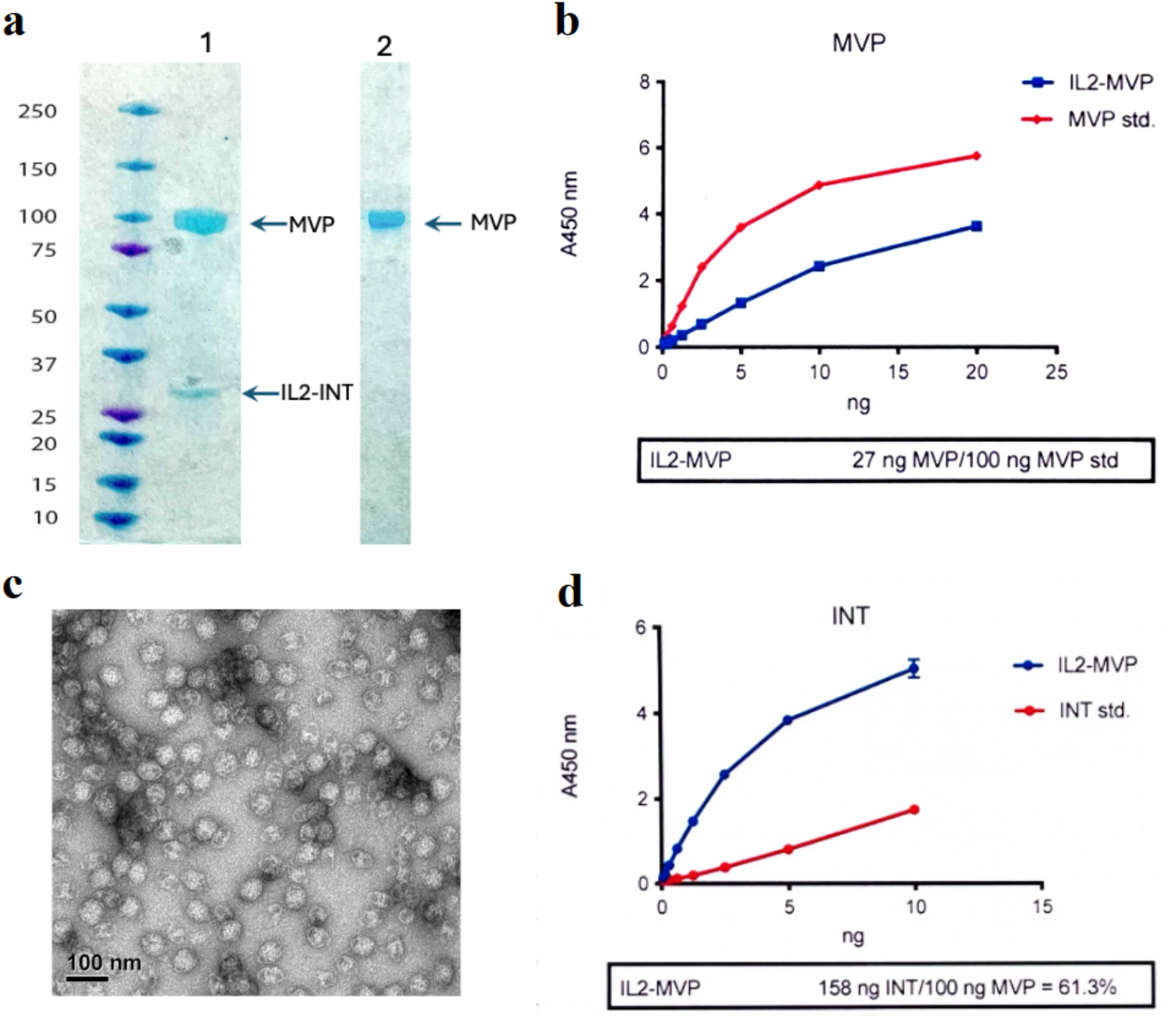
Biochemical and negative-stain EM characterization of vault–IL2 and empty vault particles. **a** SDS PAGE, Equal amounts of vault-IL2 (lane 1) and empty vaults (lane 2) were loaded on the gel and detected by Coomassie blue staining. **b** Quantitative ELISA of vault-IL2 (labelled IL2-MVP) to determine MVP concentration. **c** Negative Stain of IL2-vaults. **d** Quantitative ELISA of vault-IL2 (labelled IL2-MVP) to determine INT concentration.

**Supplementary Fig. 2.**
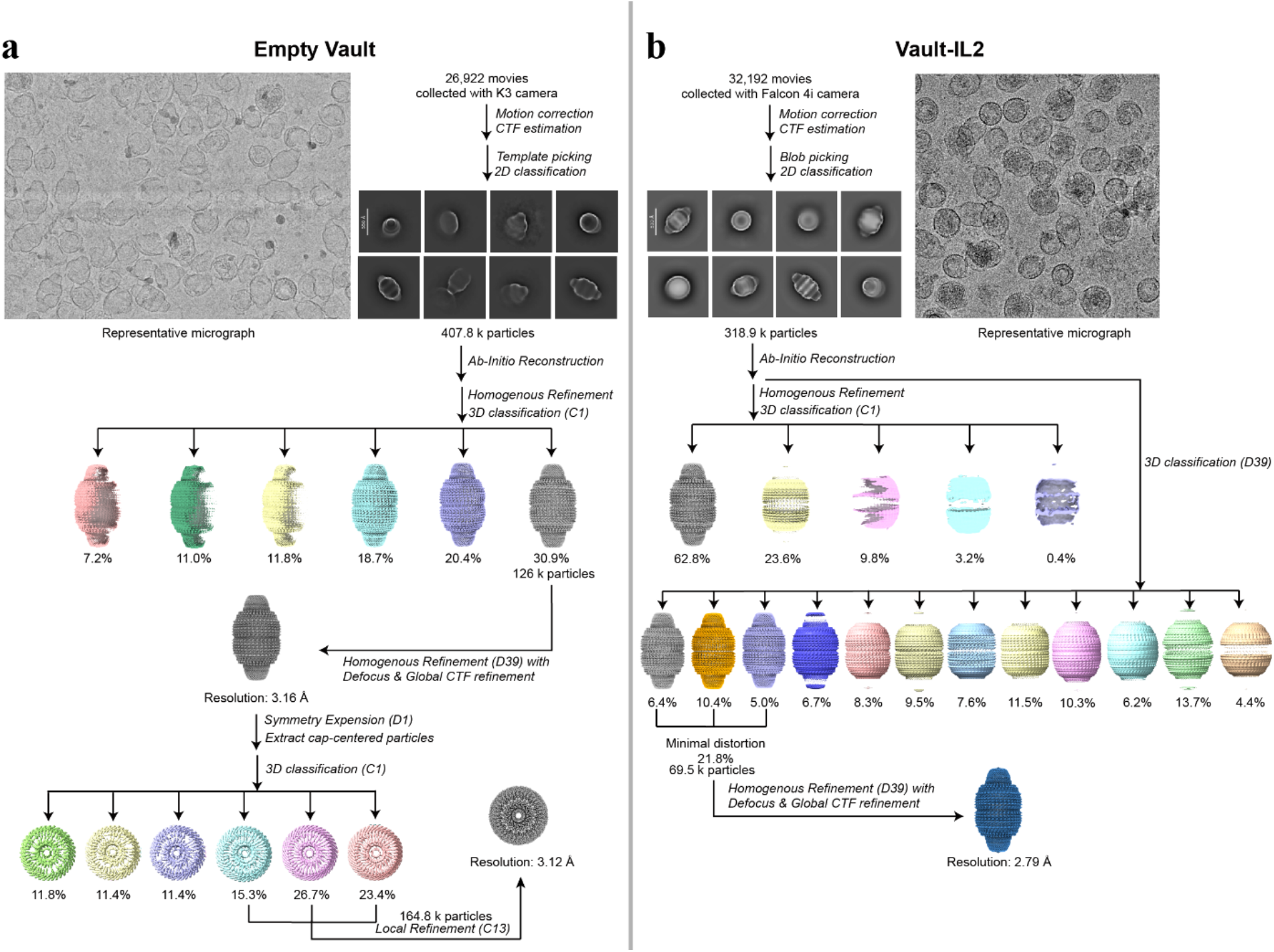
CryoEM data processing and map quality assessment of the empty vault and vault–IL2 structures. **a–b** CryoEM data processing workflow for the empty vault (**a**) and vault–IL2 (**b**) datasets.

**Supplementary Fig. 3.**
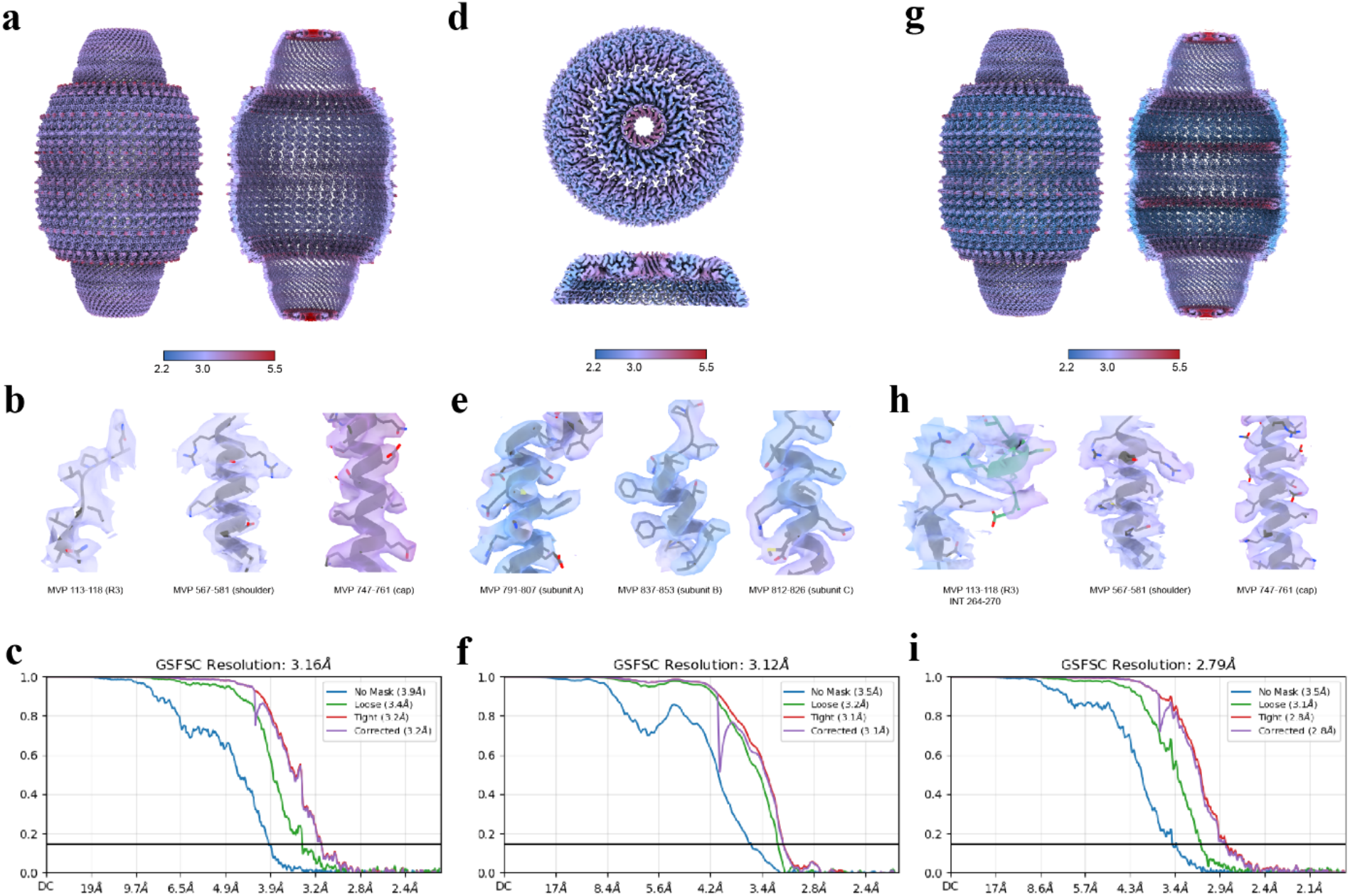
CryoEM reconstruction quality and representative model fitting. **a-c** Local resolution distributions (**a**), representative cryoEM densities and fitted atomic models (**b**) and gold-standard FSC curve (**c**) of the empty vault cryoEM reconstruction. **d-f** Local resolution distributions (**d**), representative cryo-EM densities and fitted atomic models (**e**) and gold-standard FSC curve (**f**) of the cap-focused reconstruction. **g-i** Local resolution distributions (**g**), representative cryoEM densities and fitted atomic models (**h**) and gold-standard FSC curve (**i**) of the vault-IL2 reconstruction.

**Supplementary Fig. 4.**
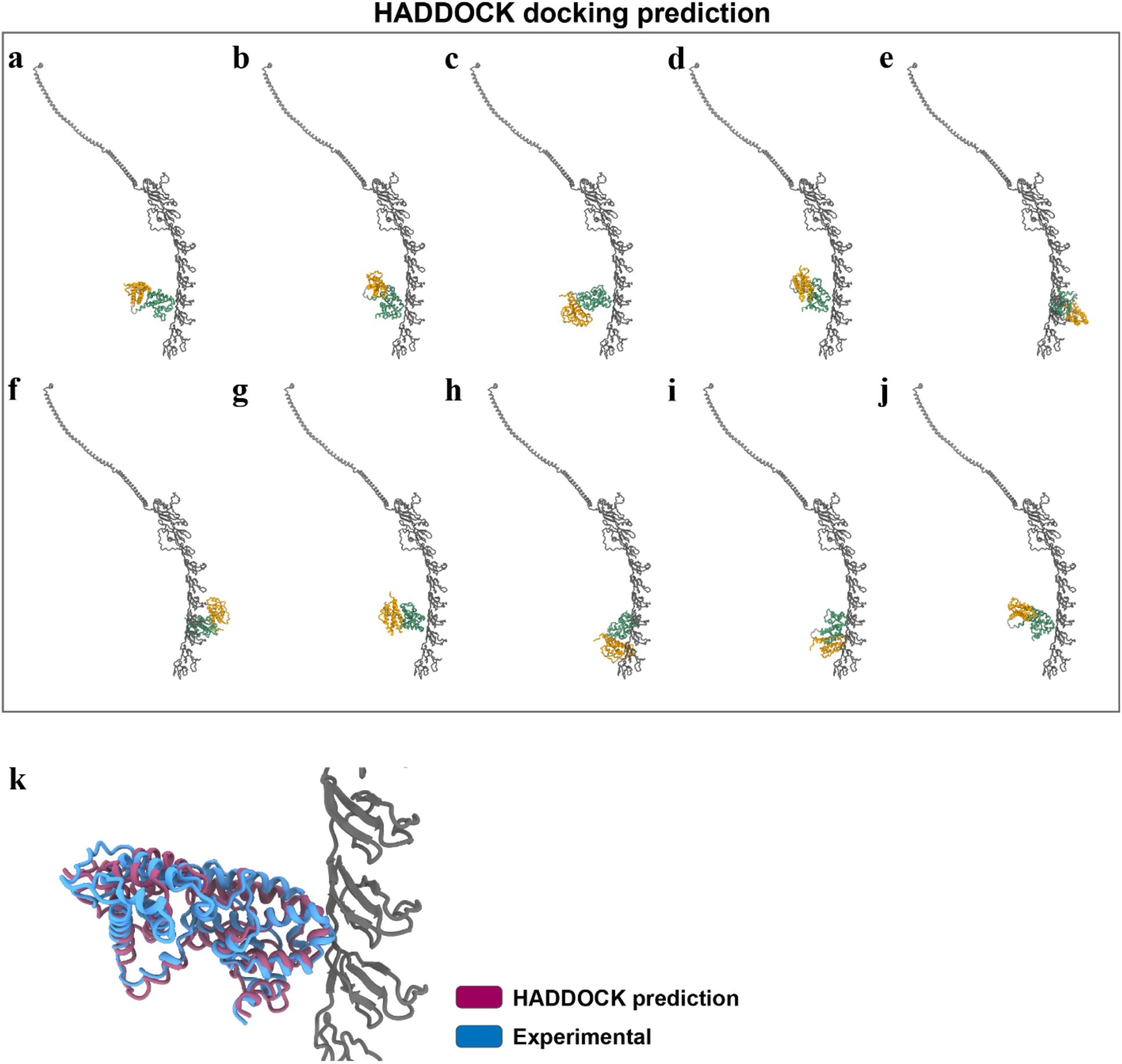
HADDOCK predictions of INT–MVP interactions. **a–j** Top ten HADDOCK docking solutions for the interaction between MVP and the AlphaFold3-predicted IL2-INT cargo, generated using predefined candidate interaction restraints on INT and the MVP R3–R4 region. **k** Structural alignment of the top-ranked HADDOCK docking model and the cryoEM-derived INT-binding model. The restraint-guided docking solutions are compatible with the experimentally observed INT–MVP binding mode at the R3– R4 region.

**Supplementary Fig. 5.**
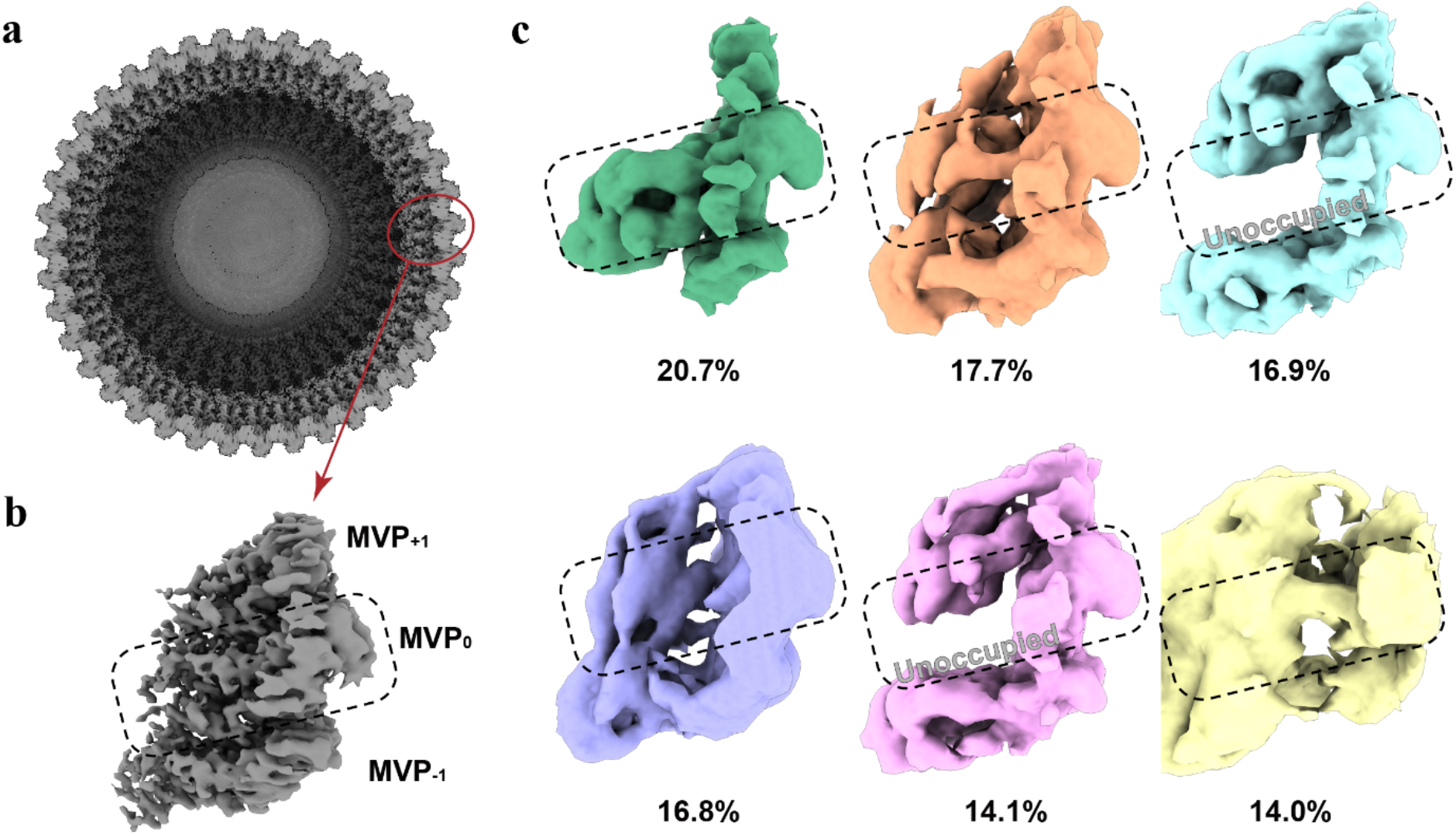
Heterogeneous occupancy at the INT-binding site. **a** Guide map of the vault–IL2 reconstruction showing the location of the cropped INT-binding region used for focused classification. **b** Reconstruction generated from the symmetry-expanded and subtracted particles. **c** 3D classification of the subtracted INT-binding region. Classes with little or no detectable cargo density were observed, indicating the presence of unoccupied binding sites. Percentages indicate the fraction of particles assigned to each class.

**Supplementary Fig. 6.**
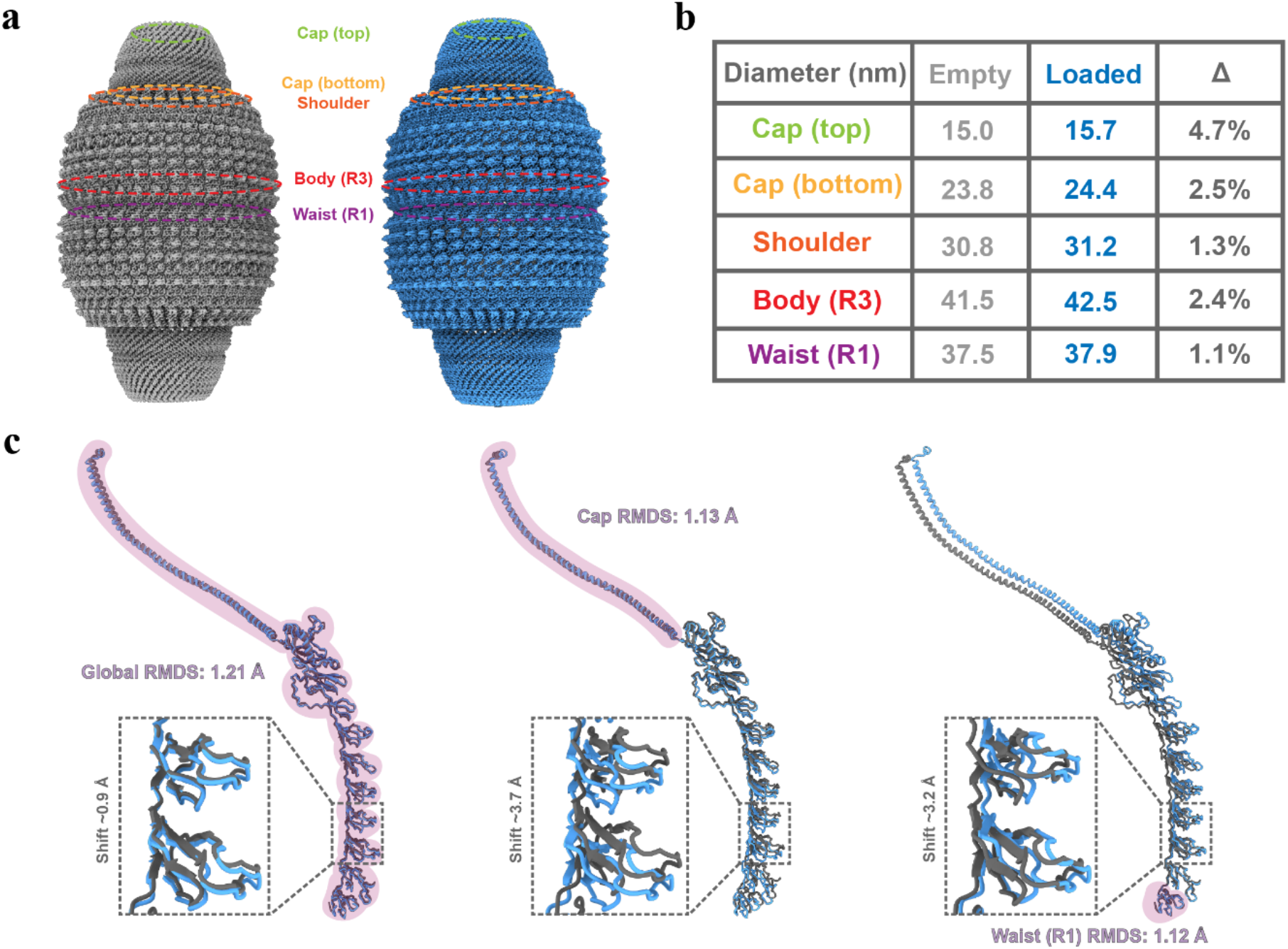
Structural comparison between the empty vault and vault–IL2 complex. **a** Superposition of empty vault (gray) and vault–IL2 (blue) structures. **b** Quantitative comparison of diameters measured at five characteristic regions (top cap, bottom cap, shoulder, body/R3, waist/R1). Cargo loading induces modest expansion, most pronounced at the cap. **c** Structural superposition of MVP monomers from empty (gray) and packaged (blue) vaults. Global alignment and alignments using cap or waist as reference are shown (pink highlights). Insets show local shifts at the R3–R4 INT-binding region (RMSD values and residue displacements indicated).

**Supplementary Fig. 7.**
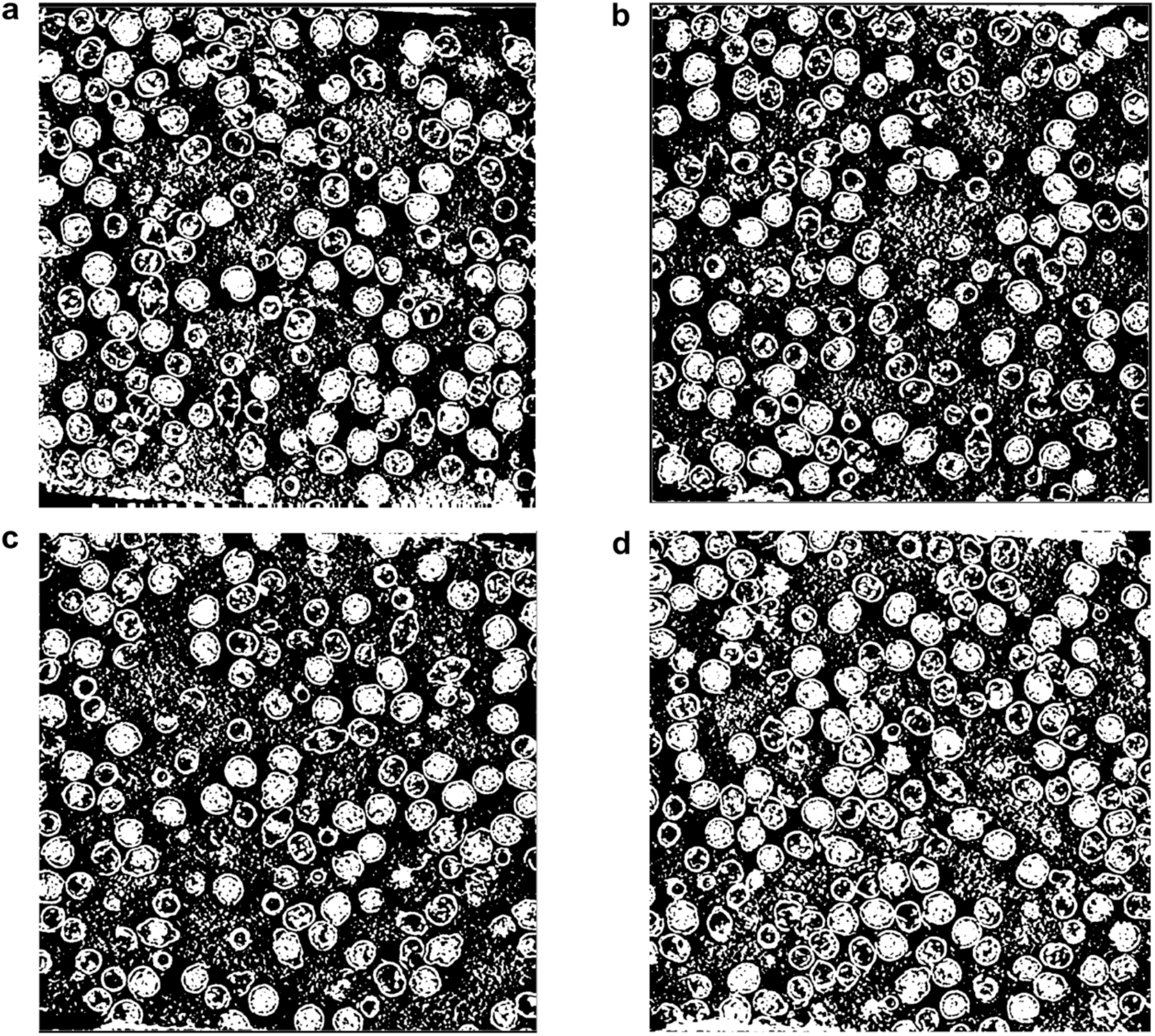
Chosen binary classification thresholds for quantitative cryo-ET analysis. **a-d** Exemplary thresholds chosen for different slices of the 4 tomograms. White pixels are considered to be above the threshold, and black pixels are considered below the threshold.

**Supplementary Fig. 8.**
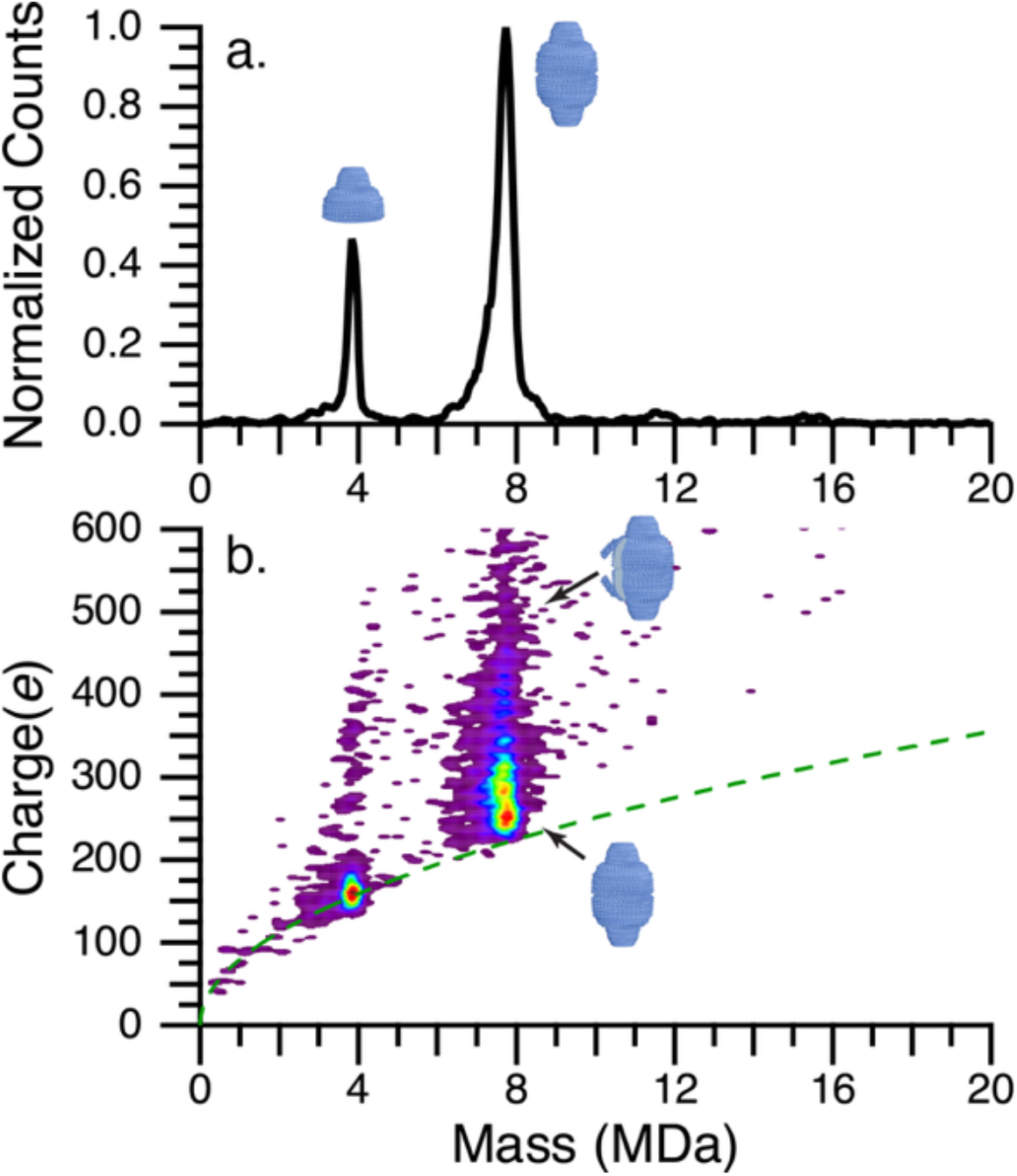
CDMS spectra of empty vaults acquired under lower *m*/*z* tuning conditions to better transmit more highly charged ions. **a.** 1D mass histogram of empty vault particles. **b.** 2D mass / charge histogram of empty vault particles. The Rayleigh limit computed for a spherical water droplet is given by the green dashed line.

**Supplementary Fig. 9.**
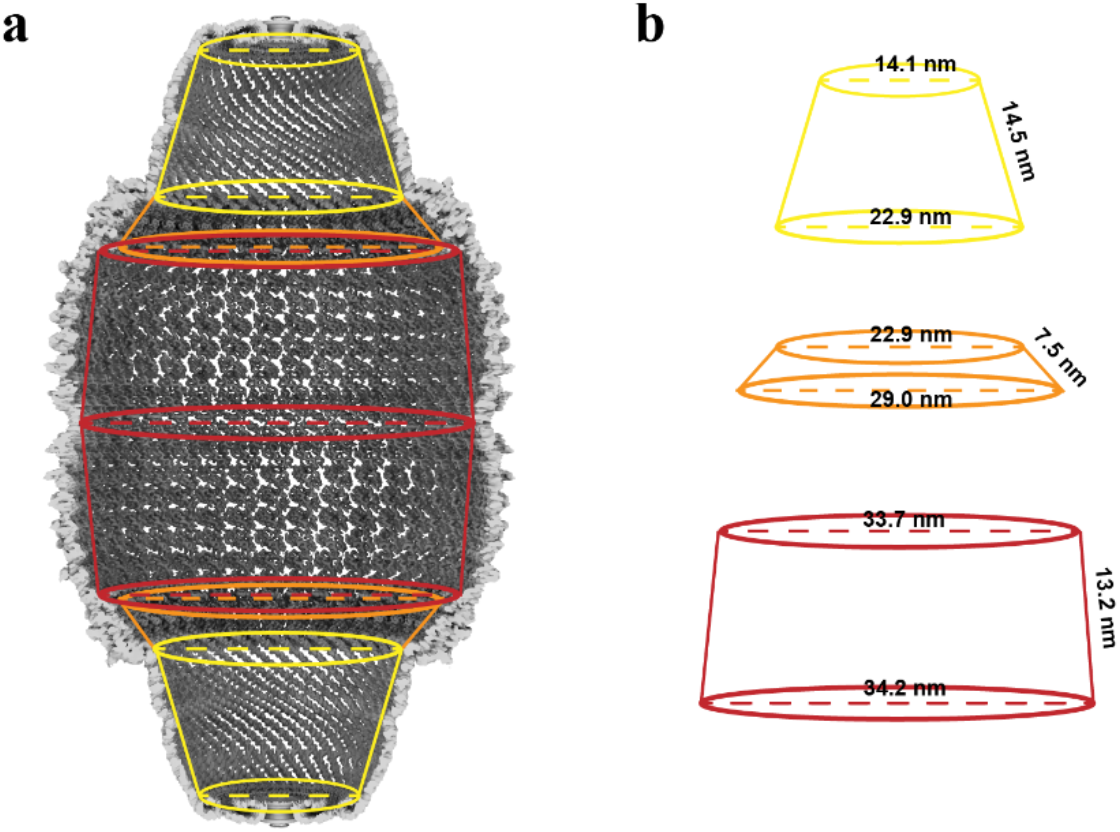
Geometric estimation of vault lumen volume. **a** Simplified frustum-based geometric approximation on the empty-vault density map. The vault lumen was approximated as a series of truncated-cone segments for volume estimation. **b** Schematic of the geometric segments and dimensions used for the volume calculation. Dashed lines indicate measured internal diameters, and side labels indicate measured slanted side lengths. The volume of the vault was approximated by dividing it into the three conical frustums shown. The volume, *V_frustum_*, of each conical frustum was computed using the following equation, where *r_1_* and *r_2_* are one half of the larger and smaller diameters, respectively, *s* is the slant length and *h* is the overall height.

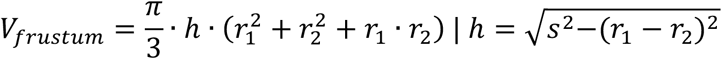 The sum of twice the volume of the three conical frustums gives an approximate volume for the vault. Using the dimensions listed, the empty vault has an internal volume of 3.87×10^−20^ liters. The maximum packaging with no internal solvent ∼960 copies if a density of 1.42 g/mL for 34.5 kDa IL2-INT is used.^29^

**Supplementary Fig. 10.**
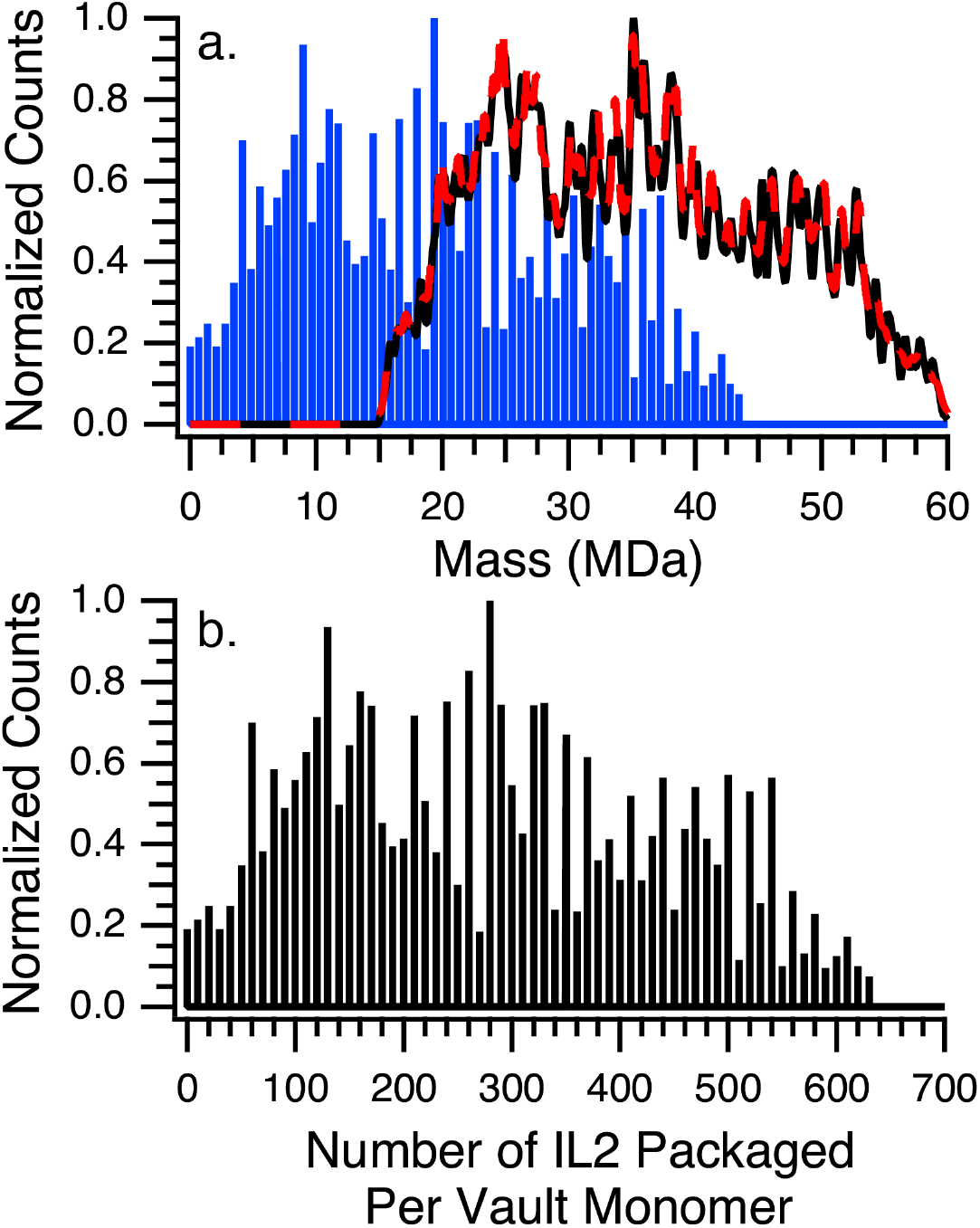
Deconvolution of IL2-INT packaging within vault dimers from Figure 3a (blue polygon). **a.** 1D mass histogram of IL2-INT packaged vault monomers (black data), least squares fit (red dashed line) and deconvolved packaging distribution (blue data). The blue lines are spaced by 20 equivalents of the IL2-INT protein (34.5 kDa). **b.** Packaging distribution from (a) where the mass axis has been converted to the number of IL2-INT copies. The population of packaged vaults that are dimers (∼22%) is comparable to that for the empty dimers (∼24%). These dimers likely exist in solution owing to the small emitter tips used which greatly reduce or eliminate the extent of non-specific aggregation that can occur due to interactions during the ESI process. The packaging distribution is broad and extends from zero to 600+ IL2-INT copies per vault monomer. The average packaging per vault for the dimers (∼281) is slightly higher than that for the monomers (∼230) suggesting that the bloated vaults may have a slightly higher propensity to form dimers.

**Supplementary Fig. 11.**
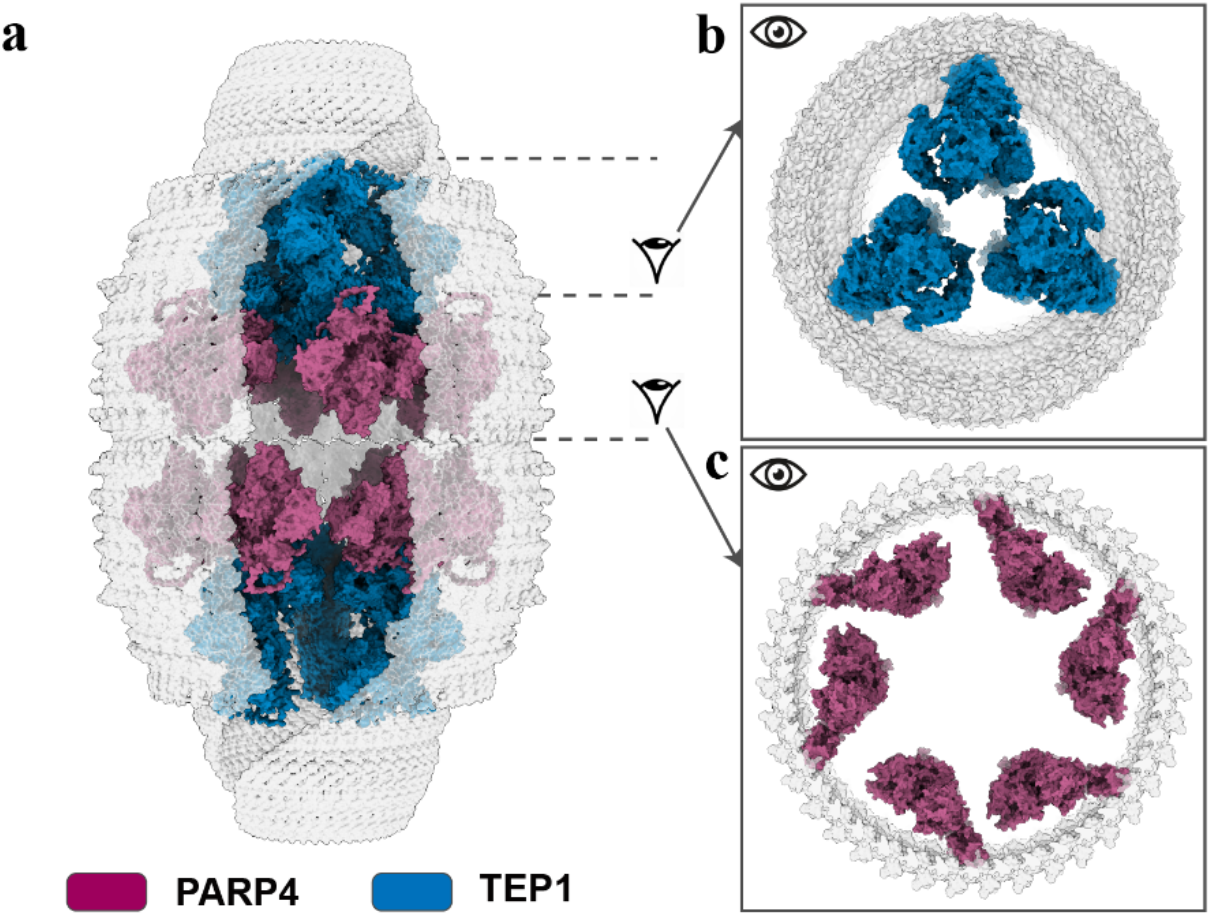
Illustrative packing model of endogenous vault-associated proteins within the vault lumen. **a** Manual steric-feasibility models of PARP4 and TEP1 positioned within the lumen. PARP4 is shown in magenta and TEP1 in blue; the vault shell is shown in transparent gray, with 10 front-facing MVP subunits hidden to expose the internal packing arrangement. The models were generated from AlphaFold3 predictions and manually positioned to estimate the maximum number that could be accommodated within the internal cavity. Long flexible and low-confident regions that are not structurally constrained were removed for visualization and packing analysis. These regions included PARP4 residues 1207–1560 and TEP1 residues 1–233 and 2466–2538. There is ample space for these regions inside the vault, but AlphaFold3 cannot predict their precise location. The PARP4 INT region was manually reoriented to align it to the INT-MVP binding location determined by our high-resolution structure. **b, c** Axial views showing representative packing arrangements of TEP1 (**b**) and PARP4 (**c**), respectively. These models are intended as illustrative steric-packing estimates rather than definitive structures or native stoichiometry models.

